# Intraspecific glucosinolate chemodiversity in *Cardamine hirsuta*

**DOI:** 10.64898/2026.11.11.765667

**Authors:** Niels Agerbirk, Karin Djendouci, Marcus A. Koch, Dominik Ziaja, Caroline Müller

## Abstract

Within some plant species, a high intraspecific chemical variation, termed chemodiversity, can be found in the composition of specialized metabolites. For Brassicales, glucosinolates are highly characteristic. Their composition is genetically determined but variation has been found within species and between organs in different species. *Cardamine hirsuta* is increasingly being used as genetic model species, but until now, little was known about its genetically-based variation in glucosinolates. Thus, we aimed to characterize glucosinolate profiles, their putative genetic variation and their chemodiversity in established accessions and wild-collected material.

We analyzed the glucosinolate profiles of leaves and roots from offspring, grown under standardized conditions, of 18 accessions collected in nine countries. Glucosinolates were quantified by HPLC-DAD and identities were confirmed by UHPLC-QToF-MS/MS. Glucosinolate composition was analyzed in biosynthetic terms, compared across samples with principal component analysis and chemodiversity was quantified as glucosinolate richness and Shannon diversity.

In total, 14 glucosinolates were confirmed across samples. Two main sources of variation among accessions were the relative levels of but-3-enyl-*versus* pent-4-enylglucosinolate and of benzylglucosinolate *versus* phenethylglucosinolate. An almost universal presence of 1-methoxyindol-3-ylmethylglucosinolate in roots but not leaves was observed, along with quantitative contrasts between leaves and roots. The glucosinolate composition varied profoundly among accessions and between leaves and roots, while offspring within accessions were mostly consistent in their profiles. Even accessions collected in close proximity produced distinct profiles, with no clear separation along latitude or longitude. Our study highlights two independent and chemically simple traits of apparently genetic nature, namely the extent of chain elongation of methionine-derived and of phenylalanine-derived glucosinolates. Their independent occurrence is a main factor shaping the glucosinolate profiles among accessions. High variation was also observed at a local scale, highlighting complex genetic variation in glucosinolate biosynthesis in the species throughout Europe. Moreover, within individuals different organs showed clearly distinct profiles, indicating organ-specific biosynthesis. The genetic basis and ecological consequences of this variation need to be explored in future studies.

## Background

Within species, polymorphisms occur in a lot of different traits. Some species show highly obvious intraspecific variation in morphological traits such as flower color or leaf architecture [1, 2], while other variation may be more hidden, such as differences in physiology, i.e. photosynthesis [3], or biochemistry of specialized (previously termed secondary) metabolites linked to plant defense [4]. Differences in mechanical or chemical defense traits within species may be stable as they can serve ecological advantages and offer associational effects [5–8]. Chemical diversity – or chemodiversity – within plant species occurs with regard to different chemical families, such as terpenes [9, 10], alkaloids [11, 12] or glucosinolates [13–15]. These intraspecific differences can also play crucial roles for humans, if the compounds differ in toxicity or their medical value [16, 17]. However, often the genetic mechanisms underlying these polymorphisms are only poorly understood (but see, e.g., [18, 19]. Moreover, the chemotypic variation, leading to the classification into chemotypes [20], can pronouncedly differ among organs within individuals. Such differences may be based on organ-specific metabolism, differential transport and distinct ecological functions [20], but in how far chemotypic variation differs among organs has been often neglected.

Glucosinolates form a fascinating chemical family that are very well studied in terms of their biosynthesis and structural diversity [21]. Their basic structure is a *S-β-*D-glucopyrano unit which is connected to an *O-*sulfated (*Z)-*thiohydroximate function, while groups of different glucosinolates are derived from distinct amino acids [21]. Each group of glucosinolates that is derived from one shared amino acid can be referred to as a biosynthetic family. However, our current understanding of glucosinolate biosynthesis relies mostly on certain glucosinolate groups for which genetic variation exists in the model plant *Arabidopsis thaliana* and in a few *Brassica* crops [22, 23], where predominantly glucosinolates derived from the amino acids tryptophan or methionine have been studied. Studies of species with other glucosinolate biosynthetic families can contribute functionally different genes for use in plant breeding or biotechnology [24, 25] and are also needed for a wider understanding of past and present glucosinolate evolution and its ecological consequences.

Plant species differing in their glucosinolate composition have been found to evoke different ecological functions, impacting the preference and performance of herbivores and pathogens. For example, populations of *Bunias orientalis* differing in fingerprints and glucosinolate profiles caused distinct performance in a polyphagous herbivore species and its parasitoid [26]. Nevertheless, the reported effects may also be due to other compounds that co-vary with the glucosinolates. Likewise, a glucosinolate polymorphism in leaves of *Barbarea vulgaris* was associated with different impacts on Lepidopteran species [27]. However, this suggested causal effect of the *B. vulgaris* glucosinolate polymorphism on insect herbivores could not be supported when the pure glucosinolates were topically applied to other Brassicaceae plants [28]. Another polymorphism in *B. vulgaris* that leads to a distinction of chemotypes was attributed to co-varying antifeedant saponins [29]. To understand the mechanisms underlying such polymorphisms, the genetic underpinnings need to be studied. Moreover, to understand potential distinct ecological effects of different accessions on interacting species, the actual natural chemodiversity should be investigated in detail. Indeed, not only individual glucosinolates but also their richness and overall diversity (e.g., calculated as Shannon diversity) may impact the defensiveness, or defense syndromes, of Brassicales [15, 30, 31].

*Cardamine hirsuta* L. is native to Europe but has been introduced to other parts of the world where it is partly invasive [32]. It has become a model species for genetic [33, 34] as well as ecological studies [32, 35]. In contrast to *A. thaliana* [36–38], *C. hirsuta* has not been extensively investigated for variation in glucosinolate profiles. In a preliminary investigation glucosinolate profiles of just two accessions were compared [39], but during the present investigation it was revealed that these included a misidentified accession of *Cardamine flexuosa* (Agerbirk and Koch, erratum in prep.).

The aim of this study was to investigate the inter- and intra-individual glucosinolate profile variation of *C. hirsuta* in a considerable number of accessions, focusing on both local variation and variation at a European scale. To exclude the impact of different environments, offspring from 18 accessions (10 from Southern Scandinavia, 8 from a wider geographic range) were grown under standardized conditions and leaves and roots of comparable developmental stage were harvested. Fourteen glucosinolates were found to constitute the majority of glucosinolates in all offspring plants, with considerable variation of profiles among accessions. The glucosinolate composition was analyzed in the framework of known glucosinolate biosynthesis genes in *A. thaliana*, as well as from an ecological perspective comparing the chemodiversity using principal component analyses and statistically analyzing the richness and Shannon diversity. We report extensive and complex variation in glucosinolate profiles in *C. hirsuta*. Because of the uniform growth conditions and uniform results for most accessions, the variation seems to have a genetic basis, but in several ways does not correspond closely to genetic variation studied in other crops or model plants. Principal component analysis revealed that the individual accessions were relatively uniform but glucosinolate profiles were quite distinct between leaves and roots. Moreover, the glucosinolate analyses highlight considerable variation on a very local scale as well as on a European scale.

## Materials and methods

### Plant growth

Seeds of different accessions (all taken from individual plants, Table 1) were sent to Bielefeld University and placed on glass beads in Petri dishes in a climate chamber with 20 °C/16 °C and 16 h light :8 h dark and 70% r.h. for germination. Seedlings were transferred into individual pots (12 cm diameter) filled with a 2:1 mixture of soil (Fruhstorfer T-type, HAWITA GmbH, Vechta, Germany): sand and fertilized with 10ml 0.1% Wuxal Super (8-8-6 + Sp, Hauert MANNA, Nürnberg, Germany) per week, starting 17 day after potting. After 62 – 66 days, ideally before plants started flowering, plant parts were harvested. Six middle-aged leaves per individual were transferred in Eppendorf tubes and directly snap-frozen in liquid nitrogen. For root harvest, soil was removed, roots cleaned by rinsing in water, gently drying with paper and also snap-frozen. This process took at maximum 20 min per plant. Plant material was kept frozen at −70 °C. Per accession, leaves and roots were taken from 3-6 individuals. If existing, further individuals per accession were kept for longer and in case they started flowering, they were used to prepare an herbarium voucher. Since not all accessions had sufficient plants growing or plants did not flower within seven months, vouchers are not available from all *C. hirsuta* plants. Vouchers are stored at Heidelberg University, Germany.

**Table 1:** Accessions, provider and code, country of origin, GPS coordinates, voucher ID for entire dried plant and ID for dried leaf material (stored in herbarium of Heidelberg University, Germany). Accessions are ordered by longitude.

| Code | Provider-code<sup>1</sup> | Country | Latitude | Longitude | Voucher ID (full plant) | Herbarium number for dried material |
|---|---|---|---|---|---|---|
| US1 | MPIPZ 7437410 | USA | 45.78333 | -122.53333 | HEID506625,<br>HEID506626 | HEID8013052 |
| GB1 | MPIPZ 7437319 | United Kingdom | 50.01647 | -5.25358 | HEID506630,<br>HEID506631 | HEID8013051 |
| ES1 | MPIPZ 7437095 | Spain | 43.43884 | -4.85157 | HEID506623,<br>HEID506624 | HEID8013050 |
| GB2 | CZ OXF-276 | United Kingdom | 51.45 | 1.15 | HEID506634,<br>HEID506635 | HEID8013043 |
| NL1 | MPIPZ 7437241 | The Netherlands | 52.15227 | 6.7033 | HEID506636,<br>HEID506637 | HEID8013049 |
| DK4 | DK-4 | Denmark | 56.064 | 9.942 |  | HEID8013046 |
| DK2 | DK-23-2a | Denmark | 55.766 | 12.477 | HEID506639,<br>HEID506640 | HEID8013045 |
| DK1 | DK-23-1a | Denmark | 55.776 | 12.482 |  | HEID8013044 |
| SE1 | SE-23-1 | Sweden | 56.46 | 12.68 | HEID506627,<br>HEID506628,<br>HEID506629 | HEID8013053 |
| SE2 | SE-23-2 | Sweden | 56.46 | 12.68 |  | HEID8013054 |
| SE3 | SE-23-3 | Sweden | 56.46 | 12.68 |  | HEID8013055 |
| SE6 | SE-23-6 | Sweden | 56.46 | 12.73 |  | HEID8013056 |
| SE4 | SE-23-4 | Sweden | 56.45 | 12.75 |  | HEID8013057 |
| SE5 | SE-23-5 | Sweden | 56.45 | 12.75 |  | HEID8013058 |
| SE7 | SE-23-7 | Sweden | 56.44 | 12.81 |  | HEID8013059 |
| DE1 | MPIPZ 7437204 | Germany | 52.53427 | 13.4124 | HEID506637,<br>HEID506638 | HEID8013048 |
| SK1 | CZ-H-5 | Slovakia | 48.17 | 17.06 | HEID506632,<br>HEID506633 | HEID8013042 |
| GR1 | MPIPZ 7437247 | Greece | 40.62117 | 23.1918 |  | HEID8013047 |

<sup>1</sup> MPIMZ: seeds from collection of Miltos Tsiantis, Max Plank Institute for Plant Breeding, Cologne, Germany (2014, 2015, 2016); CZ: samples by Martin Lysák (Masaryk University, Brno, Czech Republic; collected before 2014). Accessions from Sweden and Denmark were field collected in 2023 as seeds of one single plant each by Niels Agerbirk at the following habitat types DK1 and DK4, road sides; DK2, collector’s own garden 50 m from DK1; SE1, SE2 and SE3, coastal plain quite close to each other; SE4 and SE5, agricultural pastures quite close to each other; SE6 exposed hill with natural, open vegetation; SE7, lawn between garden and coast. DK3 was excluded from the investigation, because seeds had been pooled from three plants.

### Glucosinolate extraction and quantification

Plant material was lyophilized, dry mass taken und samples homogenized with a ball mill. About 15 mg of powder (for 9 root samples less) were extracted threefold in 1 mL 80% methanol (HPLC-grade, Fisher Scientific, Schwerte Germany), adding 2-propenylglucosinolate (Sigma Aldrich GmbH, Taufkirchen, Germany) as an internal standard at the first extractions. Extracts were homogenized, centrifuged and supernatants applied on anion exchange columns [Sephadex A25 (AppliChem, Darmstadt, Germany), in 0.5 M acetic acid buffer, pH = 5.0]. Columns were washed five times with 1 mL water and subsequently two times with 1 mL 0.02 M acetic acid buffer (pH = 5.0). Finally, *Helix pomatia* sulfatase (Sigma-Aldrich; in 0.02 M acetic acid buffer), purified according to [40], was applied to the columns, converting glucosinolates into desulfoglucosinolates. On the following day, columns were washed six times with 1 mL Millipore water, solutions dried and samples resolved in water. The desulfoglucosinolates were analyzed by HPLC (Dionex Ultimate 3000, Thermo Fisher Scientific, Waltham, MA, USA) coupled with a diode-array detector (210–370 nm), equipped with a Hypersil Gold column (150 x 3 mm, 3 µm, Thermo Fisher Part. No. 25003-153030), at 25 °C column temperature. A gradient of eluent A (Millipore water) and eluent B (acetonitrile, HPLC grade, Fisher Scientific) at a flow rate of 0.5 mL min^-1^ was used, starting at 2% B, increasing to 5% B until 6 min, to 15% B until 8 min, to 38% until 12.2 min, to 50% until 14.3 min, and to 98% B until 18.0 min, followed by a column cleaning and equilibration cycle. (Desulfo-)glucosinolates were identified by comparing their retention times and spectra to an in-house databank, established from purchased glucosinolates (Phytoplan Diehm & Neuberger, Heidelberg, Germany; Carl Roth GmbH, Karlsruhe, Germany) and desulfoglucosinolates purified by NA [39]. Prior using 2-propenylglucosinolate as internal standard, a set of representative samples (*n* = 20) from different accessions were extracted and analyzed without addition of 2-propenylglucosinolate, consistently showing no detectable peak of desulfo-2-propenylglucosinolate or any coeluting peak. For quantification, peak areas integrated at 229 nm were related to those of the internal standard, applying the response factors 1 for aliphatic glucosinolates (Val, Leu, Iso or Meth-derived), 0.95 for benzyl- and phenethylglucosinolate, 0.5 for 4-hydroxybenzylglucosinolate (4hBZ), 0.29 for indol-3-ylmethylglucosinolate, 0.20 for 1-methoxyindol-3-ylmethylglucosinolate and 0.25 for 4-methoxyindol-3-ylglucosinolate, respectively, and normalized to sample dry mass. In this way, we obtained reliable levels of 14 known glucosinolates in leaves and 13 in roots whenever levels were higher than *c.* 0.01-0.1 µmol/g dry wt., which was considered the realized range of limit of detection (LOD). Glucosinolates were only quantified by the peak recognition algorithm if the spectrum was unambiguous (this was also controlled manually); thus, the LOD differed somewhat for different glucosinolates. For verification of glucosinolate identities, aliquots (50 μL) of one sample per accession were lyophilized and sent to the laboratory in Copenhagen for further identification by UHPLC-QToF-MS/MS.

### Glucosinolate identity confirmation

The samples were dissolved in 100 μL water each and subjected to UHPLC-QToF-MS/MS analysis as in [41]. The data files were manually screened for the fourteen glucosinolates detected by HPLC-DAD by inspection of extracted ion chromatograms in the Compass DataAnalysis software (v. 6.0) (Bruker) at stringent mass accuracy (+/- 0.005 Da) of calculated *m*/*z* values for the proton adduct ([M+H]^+^) and major fragments observed with authentic references or predicted from the known fragmentation behavior of lower or higher homologs. For all desulfoglucosinolates, an additional *m*/*z* value concerned neutral loss of anhydroglucose [M+H-ahGlc]^+^ and for hydroxy derivatives an additional *m*/*z* value was [M+H-ahGlc-H_2_O]^+^. For indole desulfoglucosinolates, some additional fragments were included as previously reported [41].

Conclusive identification of the fourteen glucosinolates was obtained as phytochemical identification confidence level C, which is a high confidence usually indicated as “level 1” in mammalian metabolomics [42], and included match of *t*_R_, high resolution *m*/*z* value and MS2 with an authentic standard or established reference material analyzed in parallel. For most identified GSLs, isolated authentic dGSL standards were available [39], but for a few, desulfo derivatives of GSLs from *Arabidopsis thaliana* Col-0 and Kn-0 seeds were considered a reliable reference material [43]. The MS2 verification was based on the [M+H]+ ion in all cases except 4hBZ, in which case MS2 confirmation concerned the [M+H-ahGlc]^+^ ion.

Because 1mEt and the rarely considered *n*-butylglucosinolate (Bu) may coelute in HPLC after desulfation [43] several samples representing leaves, roots and a variety of accessions were analyzed in parallel with authentic Bu, 1mPr and 2mPr. We consistently detected 1mPr and 2mPr but not Bu in *C. hirsuta* samples (Fig. S1). This conclusion was confirmed from inspection of minor ions in MS2 (Fig. S2B). Examples of *m*/*z*, *t*_R_ and MS2 confirmation, in one selected sample each, of desulfated derivatives of all 14 identified glucosinolates are provided (Fig. S2).

### Barcoding of specimens

Per accession, remaining leaf material of individuals, which had been used for glucosinolate analysis, was sent to Heidelberg for further DNA barcoding experiments to confirm species identity. We sequenced the internal transcribed spacer regions ITS1 and ITS2 including the intervening 5.8 S rDNA from the nuclear genome, and from the plastid genome we sequenced *trn*L intron and *trn*L-*trn*F intergenic spacer regions [44]. Reaction aliquots contained 25 ng of genomic DNA, 0.5 μl of deoxyribonucleotide triphosphates /dNTPs) mix (10 mM), 1 x MangoTaq buffer, 1 μl MgCl2 (50 mM), 1 U MangoTaq DNA polymerase (Thermo Fischer, Dreieich, Germany) and 0.5 μl of each primer (10 μM), in a total volume of 25 μl. ITS region was amplified using a primer combination of ITS-M13(forward)

[TGTAAAACGACGGCCAGT-GGAAGGAGAAGTCGTAACAAGG] and ITS-M13(reverse) [CAGGAAACAGCTATGACC-GGGTAATCCCGCCTGACCTGG] with the following PCR scheme: hold at 95 °C for 3 min, 29 cycles of 95 °C for 30 s, 44 °C for 30 s, 72 °C for 1 min, followed 72 °C for 5 min, and a final hold at 4 °C. *Trn*L intron [*trn*L_M13_for: TGTAAAACGACGGCCAGT-CGAAATCGGTAGACGCTACG; *trn*L_M13_rev: CAGGAAACAGCTATGACC-GGGGATAGAGGGACTTGAAC] and *trn*LF [trnL_M13_for: TGTAAAACGACGGCCAGT-CGAAATCGGTAGACGCTACG; *trn*L_M13_rev: CAGGAAACAGCTATGACC-GGGGATAGAGGGACTTGAAC] were amplified accordingly using the following PCR scheme: hold at 95 °C for 2 min, 29 cycles of 95 °C for 30 s, 50 °C for 30 s, 72 °C for 1 min, followed 68 °C for 5 min, and a final hold at 4 °C. PCR products were purified using the QIAquick PCR Purification Kit Protocol (Qiagen). Custom Sanger sequencing was performed at EUROFINS (Ebersberg, Germany) with original PCR products and using standard M13 forward and reverse primers.

Raw sequencing data (forward and reverse) were imported into Geneious 11.1.5 [45] and processed further to assemble single fasta files removing primer information. Sequences were deposited with GenBank and respective Genbank accession codes are provided in Table S2.

For DNA sequence similarity searches the blastn option offered by NCBI (https://www.ncbi.nlm.nih.gov/) was used to run query sequence searches against the standard database, restricting the search to angiosperms and using the megablast (highly similar sequences) option with output size of n = 250. Expected threshold of BLAST hits being expected by chance, with the observed score or higher was set to 1e-6 (E value).

We used the ITS phylogenetic placement tool in *BrassiBase* (https://brassibase.cos.uni-heidelberg.de/bb2/) to search for the closest/identical sequences in a proven taxonomic context [46, 47]. The tool combines a “blast step” with a subsequent EPA (evolutionary placement algorithm) basically relying on RAxML (maximum likelihood approaches) and only needs a simple FASTA DNA file format as input and query sequence.

### Statistical analyses

All statistical analyses and plotting were conducted with R, version 4.6.1, using the packages car version 3.1.5, DHARMa version 0.5.0, dplyr version 1.2.1, emmeans version 2.0.4, ggplot2 version 4.0.3, janitor version 2.2.1, patchwork version 1.3.2, readr version 2.2.0, rstatix version 1.1.0 and vegan version 2.7.6.

Relative levels between certain biosynthetic pairs of glucosinolates, namely 3-butenylglucosinolate (Buen) and 4-pentenylglucosinolate (Peen) (studying Peen in relation to Buen+Peen) as well as benzylglucosinolate (BZ) and phenethylglucosinolate (PE) (studying PE in relation to BZ + PE) were calculated. Only for those calculations, in samples in which Peen was below the detection threshold for this glucosinolate (0.1 µmol/g d.w., in total 14 root samples), the registered zero Peen values were replaced by LOD/2 (i.e., 0.05 µmol/g d.w.) to avoid zero in the numerator, which would compromise the most conservative estimate of the value of the fraction (Fig. S3).

Due to some losses of samples during extraction, glucosinolate concentrations of 12 samples were only quantified for leaves and 5 samples for roots only. Therefore, the data used for the following statistical analyses was not fully balanced with respect to roots versus leaves. Glucosinolate compositions across samples were compared using principal component analysis (PCA) conducted for leaves and roots combined, as well as for both organs separately. To ensure normality, glucosinolate concentrations were log_2_ (x+1)-transformed and pareto-scaling was applied. A PERMANOVA with Euclidean distance was calculated for leaves and roots separately to assess the effect of accession on the glucosinolate composition.

To analyze how the inter-individual glucosinolate chemodiversity is affected by accession, linear models were fitted to the richness (number of glucosinolates per sample) and Shannon diversity of glucosinolates for the leaves and roots separately. Adequacy of the models was ensured by evaluating the goodness of fit using DHARMa plots. Statistical significance was assessed using Type II ANOVA followed by a Tukey post-hoc test.

## Results

### Wide range of biosynthetic groups of glucosinolates

Fourteen glucosinolates were identified in *C. hirsuta* leaves and 13 in roots (Figs 1, S2). These glucosinolates are derived from seven amino acids (Table 2). Their mean values ranged from less than 0.1 up to 42 µmol/g dry weight (dw) (Table 2). Some additional minor peaks were observed in HPLC-DAD chromatograms, which could be either other minor glucosinolates or other metabolites. Thus, we restrict our analysis here to the glucosinolates identified with certainty. Total glucosinolate concentrations in leaves were generally high, ranging from 40 to 210 µmol/g dw with an average of 104 µmol/g dw (Table 2; Supplement Table S1). In roots, the total glucosinolate concentrations were overall lower and ranged from 17 to 127 µmol/g dw, with an average of 57 µmol/g dw.

**Fig. 1.**
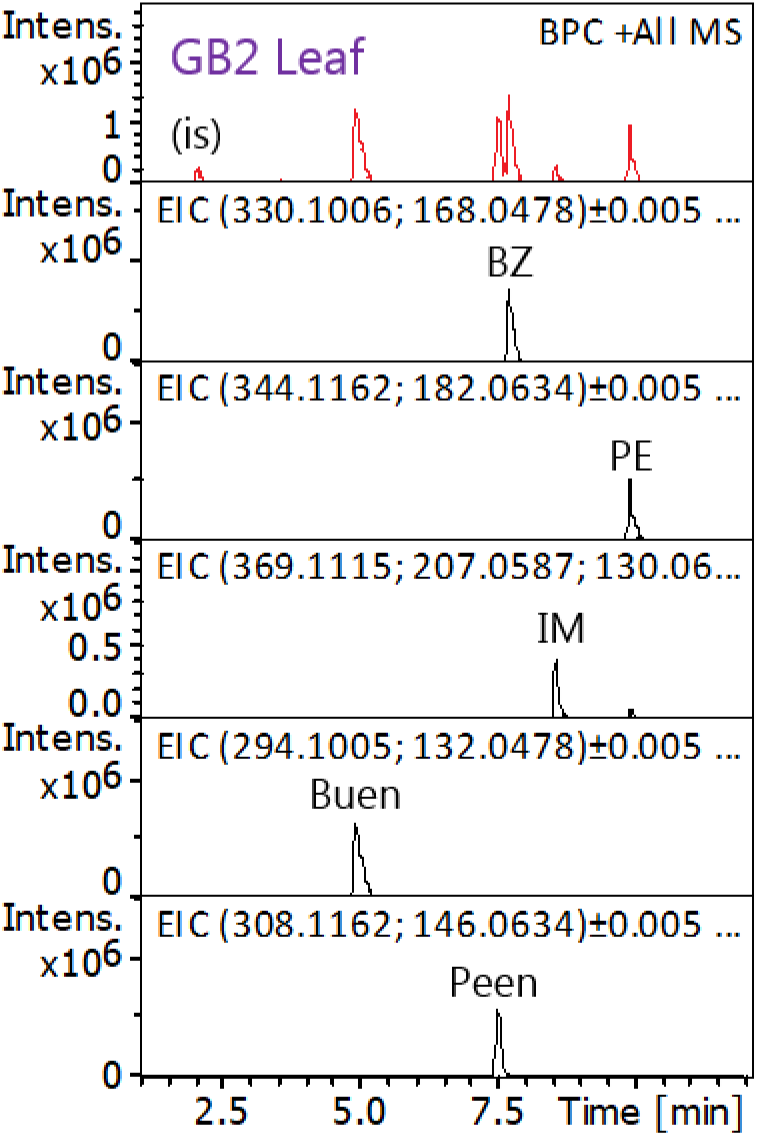
Confirmation by UHPLC-QToF-MS/MS of the identity of five major peaks from leaves of a *Cardamine hirsuta* plant from the GB2 accession. The upper (red) trace is the base peak chromatogram reflecting all major glucosinolates. The five traces below are extracted ion chromatograms each confirming the *m*/*z* values and retention times of authentic references analyzed in parallel. Confirmed peak identities, also supported by MS2 match (Fig. S2), are indicated.

**Table 2.** Mean concentrations (μmol/g dry weight) (± SD) of glucosinolates in rosette leaves (n = 79) and roots (n = 72) of *Cardamine hirsuta* offspring plants from 18 accessions as listed in Table 1. Per accession, 3-5 individual plants were analyzed. Glucosinolates (GSL) are ordered by amino acid precursor, numbers refer to [21] and retention times (*t*_R_) of HPLC-DAD are given; n.d. – not detected.

| Precursor | Glucosinolate (GSL) | Abbre-viation | No. | *t*<sub>R</sub> (min) | Leaves |  | Roots |  |
|---|---|---|---|---|---|---|---|---|
|  |  |  |  |  | **mean** | **SD** | **mean** | **SD** |
| Valine | 1-methylethylGSL | 1mEt | **56** | 6.9 | 0.228 | 0.302 | 0.011 | 0.025 |
| Leucine | 2-methylpropylGSL | 2mPr | **62** | 10.6 | 0.019 | 0.031 | n.d. |  |
| Isoleucine | 1-methylpropylGSL | 1mPr | **61** | 11.1 | 0.072 | 0.100 | 0.005 | 0.030 |
| Methionine | 4-(methylsulfinyl)butylGSL | 4mSOb | **64** | 3.4 | 0.124 | 0.212 | 0.062 | 0.109 |
|  | 5-(methylsulfinyl)propylGSL | 5mSOp | **72** | 6.1 | 0.158 | 0.339 | 0.116 | 0.221 |
|  | 6-(methylsulfinyl)hexylGSL | 6mSOh | **67** | 10.2 | 0.023 | 0.038 | 0.016 | 0.037 |
|  | but-3-enylGSL | Buen | **12** | 8.7 | 40.838 | 30.510 | 4.310 | 5.893 |
|  | pent-4-enylGSL | Peen | **101** | 11.8 | 10.314 | 9.232 | 1.629 | 2.294 |
| Phenylalanine | benzylGSL | BZ | **11** | 12.0 | 42.483 | 28.083 | 37.352 | 25.033 |
|  | phenethylGSL | PE | **105** | 13.2 | 6.459 | 6.418 | 7.520 | 9.401 |
| Tyrosine | 4-hydroxybenzylGSL | 4hBZ | **23** | 7.3 | 0.036 | 0.032 | 0.018 | 0.016 |
| Tryptophan | indol-3-ylmethylGSL | IM | **43** | 12.5 | 2.781 | 1.068 | 2.312 | 0.766 |
|  | 1-methoxyindol-3-ylmethylGSL | 1moIM | **47** | 14.0 | 0.039 | 0.188 | 3.838 | 1.366 |
|  | 4-methoxyindol-3-ylmethylGSL | 4moIM | **48** | 13.0 | 0.167 | 0.134 | 0.248 | 0.547 |
| TOTAL |  |  |  |  | **103.74** | **33.98** | **57.44** | **26.40** |

Four glucosinolates were dominant in some but not all plants: two methionine-derived glucosinolates, 3-butenylglucosinolate (Buen) and 4-pentenylglucosinolate (Peen), and two phenylalanine-derived glucosinolates, benzylglucosinolate (BZ) and phenethylglucosinolate (PE) (Fig. 2). These pairs differ chemically in the same way, with the latter having one more CH_2_ group than the former (Fig. 2), thus being pairs of homologs. Since the Buen-Peen and BZ-PE variations appeared to be a major variation in the dataset, and since individual leaves varied in total levels, we inspected the relative levels in each biosynthetic pair. Offspring from each mother plant tended to be similar in relative levels, although some variation was seen within six accessions, US1, GR1 and the offspring of four mother plants sampled in Sweden in relatively close proximity (SE1, SE2, SE3, SE5). Despite the occasional variation within siblings, there was a tendency for relative levels of Peen (in relation to Buen + Peen) to be either low (0.9-3.9 %), intermediate (17.6-20.9 %), semi-high in SE3 (26.5-31.9) or high (33.7-45.7%) in leaves (Fig. 3A). Similar groups were observed for root samples, but the minor contrast between semi-high and high groups was less obvious, suggesting fundamentally three chemotype groups with low, intermediate and high Peen (Fig. 3B).

**Fig. 2.**
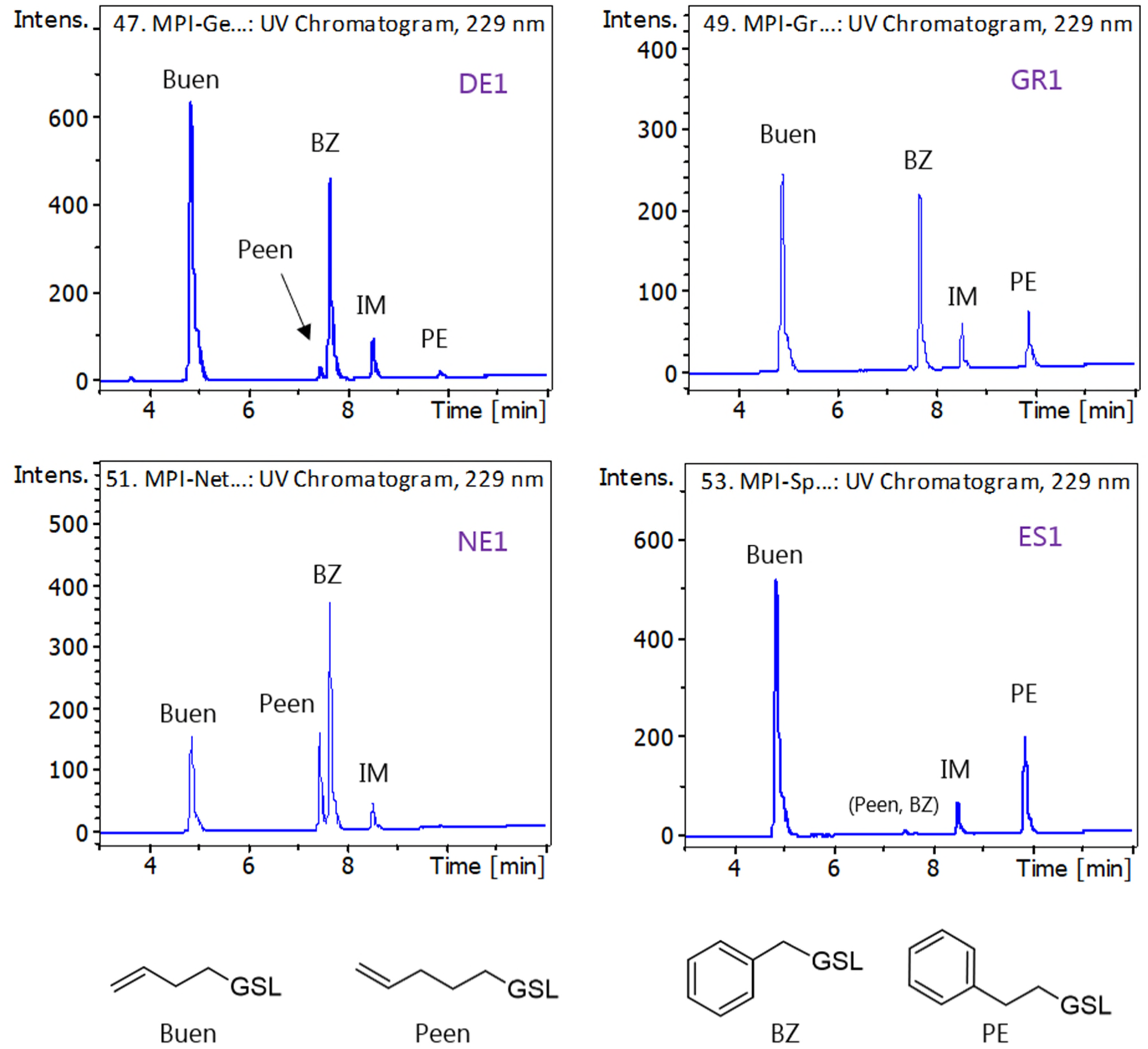
Representative HPLC chromatograms of four *Cardamine hirsuta* accessions. The accessions are named after their country of origin (for exact origins see Table 1). Structures of four glucosinolates differing in abundance among accessions are shown below the chromatograms (for glucosinolate codes see Table 2).

**Fig. 3.**
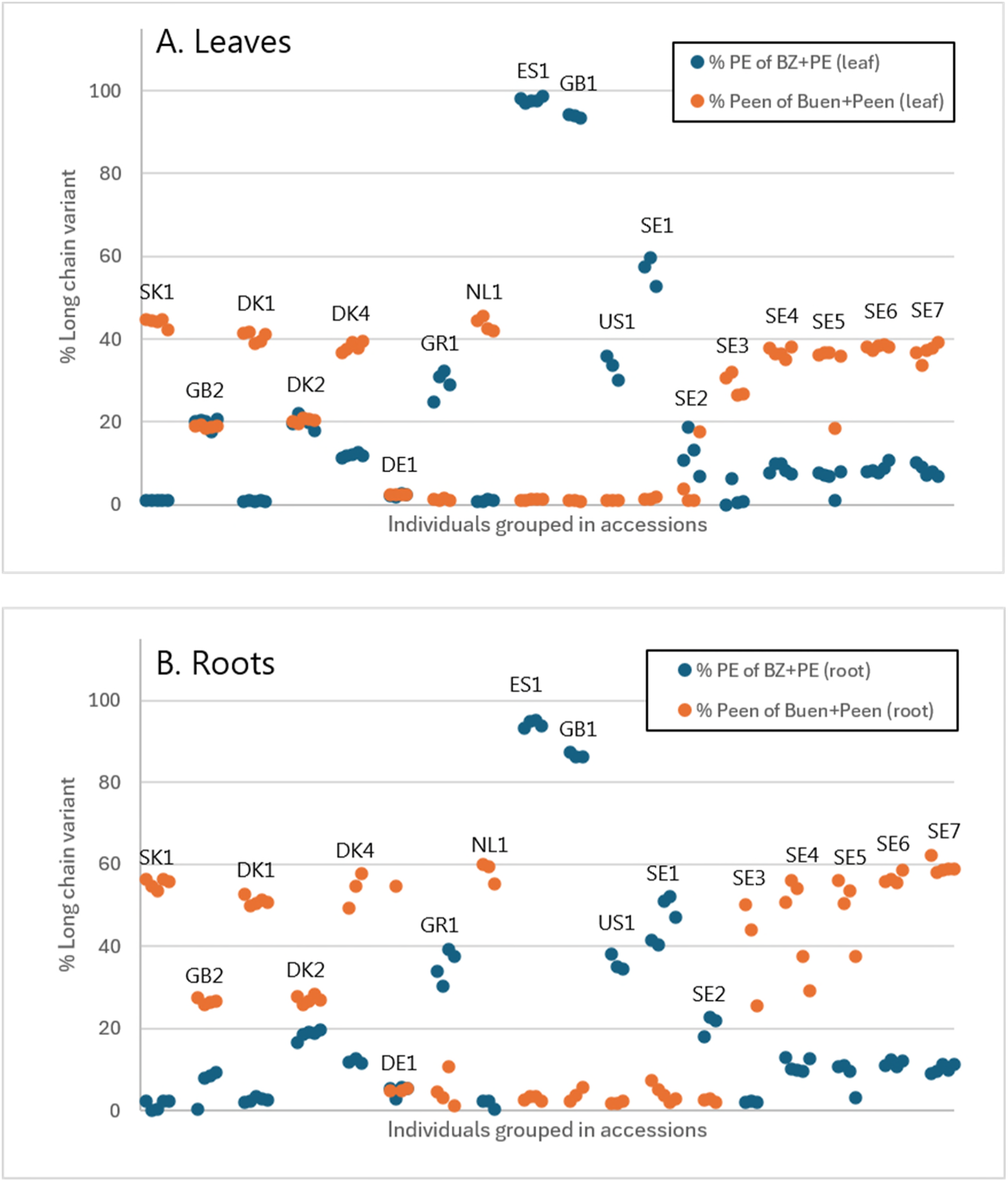
Levels of benzylglucosinolate (BZ) relative to the sum of BZ and phenethylglucosinolate (PE), and of pent-4-enylglucosinolate (Peen) relative to the sum of but-3-enylglucosinolate (Buen) and Peen, in the leaves of each of 79 *Cardamine hirsuta* plants and roots of 72 plants. Plants that were offspring of the same mother are shown consecutively. In 14 root samples, Peen was not detected but using zero for calculation of biosynthetic index appeared misleading; for this reason, the Peen level was assumed to be LOD/2 (=0.05 µmol/g) in the calculation of % Peen for those 14 plants, mainly affecting the index of one plant among SE3 (Fig. S3). For accession code see Table 1, for glucosinolate codes Table 2. Between 3 and 5 offspring were analyzed per accession.

Relative levels of PE (in relation to BZ + PE) showed a less simple pattern, but approximately five or six groups could be suggested, ranging from very low relative levels of PE (c. 1%) to very high relative levels of PE (93.5-98.5 %) in leaves (Fig. 3A). The pattern observed for roots was similar, although absolute values differed slightly (Fig. 3B).

### Differences between leaves and roots

In a PCA, glucosinolate profiles of leaves and roots were fully separated along PC1, explaining 39.8% of the variation of the data set (Fig. 4A). The leaves and roots showed slightly different abundances in several glucosinolates. 2-Methylpropylglucosinolate (2mPr) was only detectable in very low amounts in few leaf samples and not at all detectable in the roots. In contrast, roots showed consistently high concentrations in one substituted tryptophan-derived glucosinolate, 1-methoxyindol-3-ylmethylglucosinolate (1moIM), while leaves had only low amounts of this glucosinolate (Table 2). The unsubstituted tryptophan-derived glucosinolate indol-3-ylmethylGSL (IM) was consistently detected at moderate levels in both, leaves and roots (Table 2, Fig. 1). Buen and Peen were more dominant in the leaves than in the roots (Fig. 4A).

**Fig. 4.**
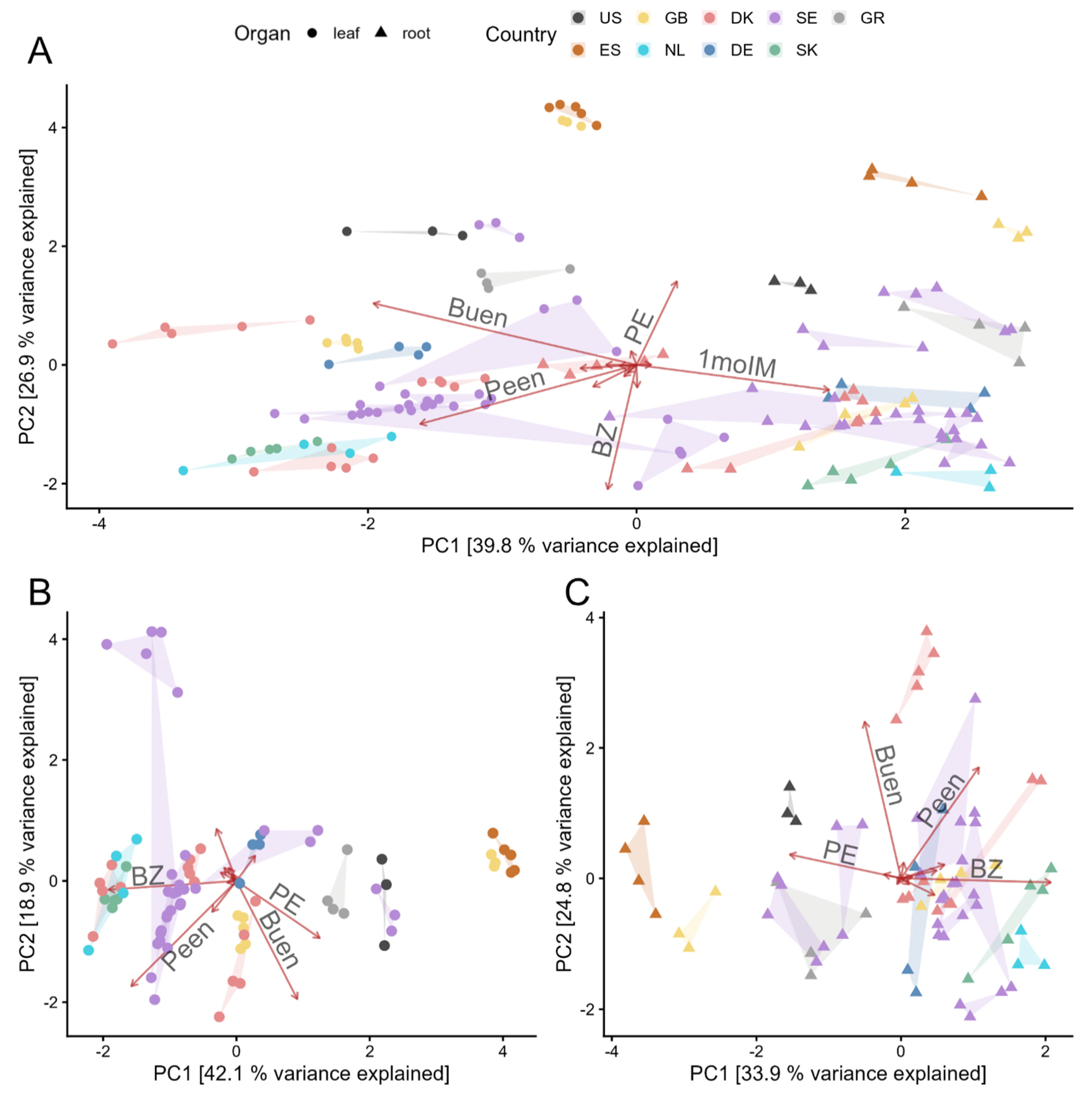
Principal component analysis of glucosinolate composition of leaves and roots together (**A**) and separately for leaves (**B**) and roots (**C**) of *Cardamine hirsuta* plants of 18 accessions. A significant (*P* < 0.001 for B and C) accession effect on the glucosinolate composition was for leaves and roots when tested separately using a PERMANOVA. Between 3 and 5 offspring were analyzed per accession.

### Differences among accessions

The glucosinolate profiles were mostly consistent within accessions, but differed strongly among accessions, being separated mainly among PC2, explaining 26.9% of the variation (Fig. 4A). Profiles of accessions differed significantly within both leaves (PERMANOVA, *F* = 22.29, *P* < 0.001; Fig. 4B) and roots (*F* = 11.07, *P* < 0.001; Fig. 4C) when tested separately. Offspring of one accession from Great Britain (GB1) and the Spanish accession (ES1) were similar but most distinct from the other accessions, both being characterized by higher levels of PE in both leaves and roots. Offspring of the three accessions from Denmark (DK1, DK2, DK4) as well as the seven accessions from Sweden (SE1 – SE7) were also mostly clearly separated from another, although seeds were collected from mother plants within a distance of 50 m for two of the Danish accessions and within an area of approximately 4 to 12 km for the seven Swedish accessions, with three contrasting accessions from the same locality (approximate distances 50-200 m between SE1, SE2 and SE3). Even those three mother plants from the same habitat produced markedly varying offspring (Fig. S3). Offspring of the accessions from the Netherlands (NL1) and Slovakia (SK1) clustered closer together, being mainly characterized by high levels of BZ in both leaves and roots. Overall, there was no clear separation across latitude or longitude.

### Chemodiversity among accessions and between organs

The number of identified glucosinolates (richness) differed significantly among accessions (leaves: *df* = 17, *F* = 9.00, *P* < 0.001; roots: *df* = 17, *F* = 7.61, *P* < 0.001), being highest in leaves of the accessions NL1, SE3 and SK1 and on average lowest in leaves of ES1 (Fig. 5A). For the roots, the richness was highest in DK2, while it was lowest in GB1 (Fig. 5B). It should be noticed, that more glucosinolates were found by UHPLC-MS of a few selected samples within the accessions, but due to low concentrations they could not be identified and quantified with certainty in the HPLC-DAD data used for the present analysis. The Shannon diversity also differed significantly among accessions (leaves: *df* = 17, *F* = 42.46, *P* < 0.001; roots: *df* = 17, *F* = 26.57, *P* < 0.001), being highest in leaves of GB2 and DK4 and particularly low in the one accession from USA (US1; Fig. 5C). In contrast, in the roots the Shannon diversity was very high in the US1 accession and in DK2, while it was lowest in NL1 and two Swedish populations (SE3, SE7; Fig. 5D).

**Fig. 5.**
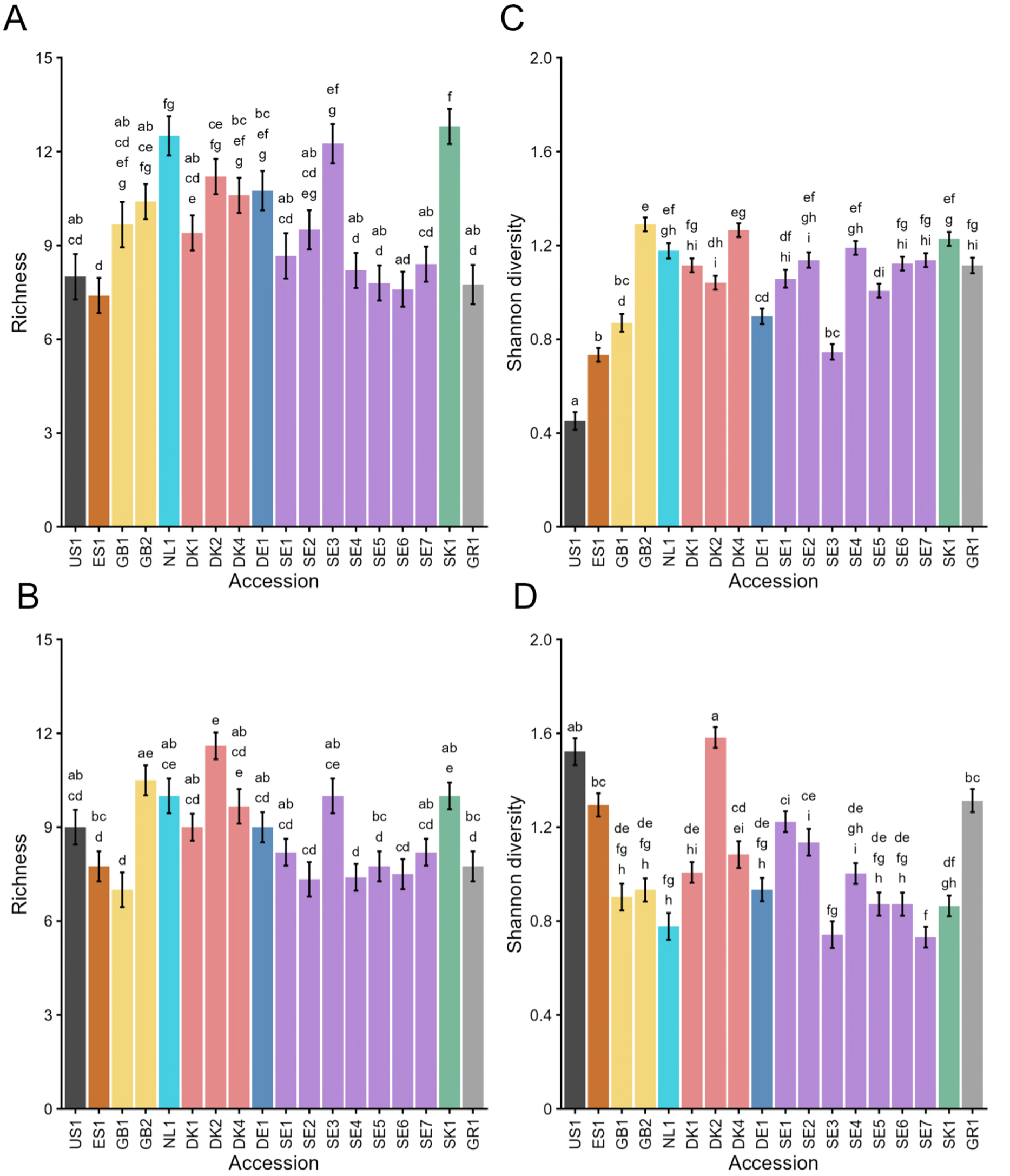
Emmeans ± standard error of richness (**A**, **B**) and Shannon diversity (**C**, **D**) of glucosinolate composition of leaves (**A**, **C**) and roots (**B**, **D**) of *Cardamine hirsuta* plants. Compact letter display is a result of a significant (*P* < 0.001 for A, B, C and D) accession effect tested for leaves and roots separately using a Type II ANOVA followed by a Tukey post-hoc test (*P* < 0.05). Between 3 and 5 offspring were analyzed per accession.

### Genetic identity

We observed very little overall genetic diversity across the three marker regions sequenced (ITS, *trn*L, *trn*LF). For ITS gene diversity *π* was calculated with 0.18%. Gene diversity for plastid *trn*L and *trn*LF were calculated with *π* = 0.4% and *π* = 1.1%, respectively. GenBank searches showed that all sequences matched a *Cardamine hirsuta* target sequence with 99.5–100% identity across the full sequence length. Among plastid reference sequences curated at GenBank, there are two entire plastome sequences from *Cardamine hirsuta* (OZ156427, NC049607). Phylogenetic placement of ITS sequences in *BrassiBase* revealed a perfect match with all curated *C. hirsuta* sequences (n = 60) referenced therein and form a monophyletic clade.

## Discussion

Overall, we identified a wide range of families of glucosinolates, representing biosynthesis from seven precursor amino acids, in *Cardamine hirsuta* collected from 18 accessions and grown under standardized conditions. Thus, the detected differences are mainly driven by genetic differences in this truly diploid species [48, 49]. The pronounced differences in glucosinolate profiles among the accessions suggest various underlying genetic principles: two major glucosinolates were Buen and Peen, whereby low levels of 4mSOb and 5mSOp would appear to be biosynthetic intermediates. These glucosinolates share a common biosynthesis in *A. thaliana* and *Brassica* spp. (Fig. 6), and chain length polymorphism among this group of glucosinolates is frequent and of considerable ecological importance [50]. However, in other genera the usual polymorphism concerns either one or two chain elongations in the biosynthesis, while Buen and Peen would require two and three chain elongations. If the diversity in *C. hirsuta* fundamentally represents mainly three levels of Peen relative to the sum of Buen and Peen, it would fit a model with just two alleles combined in heterozygotes (with intermediate levels) and two types of homozygotes (with high or low levels). Such polymorphism in *A. thaliana* concerns the MAM gene [50], with MAM2 favoring mainly one chain elongation and MAM1 favoring mainly two chain elongations. The present results are compatible with a previously unknown functional type of MAM gene in *C. hirsuta*, favoring mainly three chain elongations to synthesize 5mSOp and Peen, as well as a MAM1-like gene resulting in 4mSOb and Buen.

**Fig. 6.**
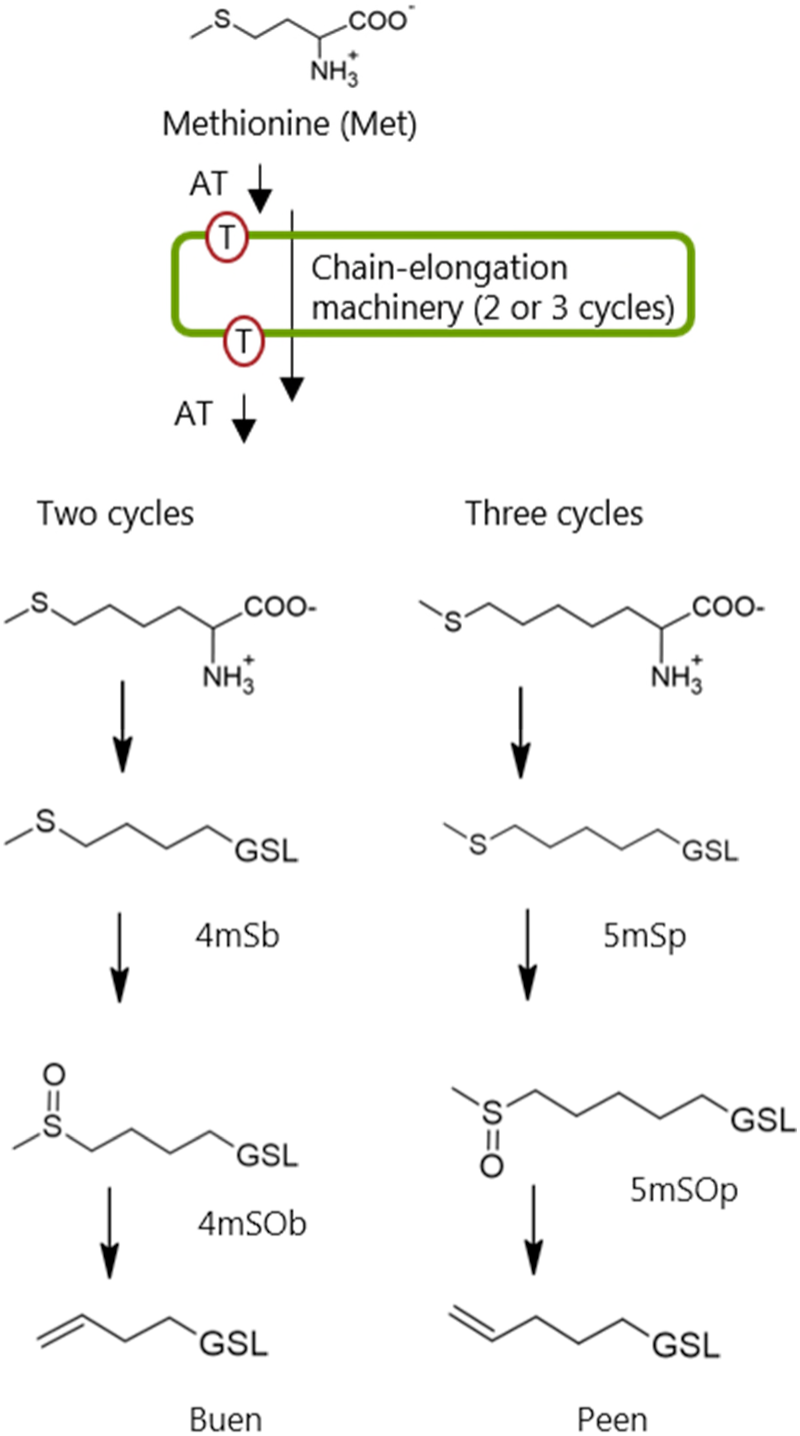
Suggested biosynthesis of methionine-derived glucosinolates in *Cardamine hirsuta*, with the Buen-Peen polymorphism explained by either two or three rounds of chain elongation. The biosynthesis follows the known biosynthesis in *Arabidopsis thaliana*, although in this species the dominating products are usually due to either one or two rounds of chain elongation. For glucosinolate abbreviations see Table 2.

An interpretation of the more complex pattern of PE relative to the sum of BZ and PE is less obvious. First, the multiple apparent phenotypes may suggest a more complex genetic background than for the Peen-Buen trait. Second, a similar polymorphism has to our knowledge never been genetically dissected. In *A. thaliana*, BZ as well as PE levels are very low, with BZ being a product of two separate and very low-expression CYP genes (CYP79A2 and CYP79C) channeling phenylalanine directly into the glucosinolate core structure biosynthesis [51, 52]. In contrast, PE in *A. thaliana* depends on one chain elongation by MAM1 of phenylalanine followed by PE biosynthesis initiated as a side product of CYP79F that mainly channels chain elongated homologs of methionine into the glucosinolate core structure biosynthesis. Hence, in the *A. thaliana* genetic framework, a chain elongation polymorphism affecting PE would be expected [43] to be correlated with chain elongation of methionine (Fig. 6), but the polymorphisms in *C. hirsuta* were clearly independent (Fig. 3).

The independent occurrence of high levels of Peen and PE relative to the lower homologs (Buen and BZ, respectively) in plants grown at uniform conditions suggests the genetic background of each variation to be independent. This independent occurrence of each kind of phenotype seems to be a major factor for the overall variation among the accessions. The considerable variation in glucosinolate profiles in both South Scandinavian areas sampled for this investigation and among the wider collection of European accessions is compatible with a stable complex genetic variation across the geographic distribution of *C. hirsuta*. This contrasts with the polymorphism found for aromatic glucosinolates in *Barbarea vulgaris* that showed an East-West organization in Europe with low local variation except at the overlap of chemotypes [53]. The current picture for *C. hirsuta* would suggest natural selection favoring no single chemotype and hence populations growing geographically mixed [13] but possibly favoring being different from plants growing close by, as is proposed by the plant community variability hypothesis [54].

In a review of glucosinolate structural variation in the Brassicaceae tribe Cardamineae, only few well-characterized intraspecific polymorphisms were revealed, such as in the sexually reproducing *Barbarea vulgaris*, in the partially apomictic *Cardamine pratensis* species complex and among wild and cultivated *Nasturtium officinale* [55]. Variation in leaf glucosinolate profiles in further Cardamineae species has also been recently described among wild populations of *Cardamine pratensis*, *C. armara* and *C. impatiens* [15]. The present detailed analysis of *C. hirsuta* at European scale, as well as of one North American accession, adds another highly polymorphic species with regard to glucosinolate structural diversity to this tribe, which supports in-tribe evolution to use and have used this kind of raw material in the continuous evolution of specialized metabolism variation. Geographic isolation of populations differing in glucosinolate profiles, followed by different selection pressures in the individuals’ specific environments, would be sufficient to result in intraspecific glucosinolate variation [53], without the need for polyploidization or gene duplication often proposed to be key factors in glucosinolate diversity evolution [56]. Distinct glucosinolate chemotypes have been also found in other tribes of the Brassicaceae, such as the Camelineae (i.e. *Arabidopsis thaliana* and *A. halleri*) [14, 22], Buniadeae (i.e. *Bunias orientalis*) [31] and Lepidieae (i.e. *Lepidium densiflorum* with contrasting glucosinolate profile among accessions [57, 58], while representative species of four other tribes collected in the wild, including the species *Lepidium draba*, *Lunaria rediviva*, *Hesperis matronalis* and *Descurainia Sophia*, showed only little variation [15]. This once again underscores that the Brassicaceae are an interesting family for studying correlations among metabolic and genetic variation, the combination of which reveals new insights.

Glucosinolate structural variation in the tribe Cardamineae so far seems to have evolved independently of chromosome number polymorphism [55]. However, the tribe Cardamineae is among those species-rich Brassicaceae clades with a high number of young polyploids (c. 60%; [59, 60]) and such polyploidization is considered as a major driver of diversification for the tribe Cardamineae and in particular for the genus *Cardamine* [61]. The complex and considerable existing variation in diploid *C. hirsuta* described here provides and alternative source of evolutionary diversification by Darwinian natural selection among intraspecific variation. A promising aspect of *C. hirsuta* is its status as a well-established genetic model plant [34, 49, 62], which may enable a genetic dissection of the kinds of polymorphism described here, and subsequent studies of ecological aspects in the field.

An apparently stable co-existence of different chemotypes within close proximity is also known for species characterized by other specialized metabolites, such as *Tanacetum vulgare* (Asteraceae) showing multiple leaf terpene chemotypes within populations [63]. A high diversity in monoterpenes among individuals of this species may be realized by unlinked enzyme loci, expression variation in alleles, presence-absence variation and *de novo* enzyme evolution [19]. In general, a high variation in chemical composition may enable chemodivers species to adapt well to different environmental conditions [5, 54, 64], with chemotypes of the same species showing, for example, different attractiveness to mutualists and antagonists [65–67]. Interestingly, herbivores in turn may also rapidly evolve adaptations to different chemotype profiles; recent work on different species of Lepidoptera feeding on Brassicaceae with distinct chemotypes revealed a high diversification in glucosinolate sulfatases to deal with glucosinolate-based defenses [68].

Along with the differences in chemotypes shaped by the Peen-Buen and BZ-PE polymorphism, also the richness of glucosinolates differed significantly among accessions of *C. hirsuta* in both leaves and roots. Overall, the glucosinolate richness was likely underestimated in the samples, as some additional putative glucosinolates were found in the samples but not yet identified with certainty and as some glucosinolates may have been in some samples below the detection limit. Differences in abiotic conditions have been found to drive the metabolite richness in willows [69] and may also have shaped the glucosinolate profiles of the accessions of *C. hirsuta* over generations. Since all plants were grown under the same conditions, these differences must be attributable to earlier selection. A distinct richness of specialized metabolites within a chemical family may impact herbivore or pollinator visits, as evidenced by correlations found in other study systems [65, 70, 71]. Within the accessions of *C. hirsuta*, the glucosinolate profiles clustered together and the richness and Shannon diversity were overall rather consistent within leaves and roots, which hints to a stable biochemical pathway among siblings. The pronounced differences in these traits even in accessions growing in close proximity to each other, such as the Swedish and Danish accessions, indicate that chemodiversity can be high on a small scale even in a comparably conserved group of specialized metabolites, such as the glucosinolates.

Apart from differences in glucosinolate composition between *C. hirsuta* accessions, we also found a pronounced difference between the glucosinolate composition of leaves and roots. One glucosinolate, 2-methylpropylglucosinolate, only occurred in leaves but in very low concentrations; it may have been simply below the detection limit in roots. In contrast, 1moIM was present in both organs, but only in traces in leaves, while in substantial concentrations in roots. This glucosinolate has also been found to be mainly root-associated in other species such as *A. thaliana* [72] and *B. vulgaris* [73]. The biological function of 1moIM includes being a precursor of indole phytoalexins [74], which may be needed to a higher extent in roots than in leaves. Overall, leaves also showed much higher total glucosinolate concentrations than the roots. The root values may have been underestimated to some extent, as cleaning of roots may have caused some losses due to an activation of myrosinases. Qualitative as well as quantitative differences in glucosinolate profiles between leaves and roots, as well as other organs, have also been revealed in other Brassicaceae [13, 75]. These may be due to different selection pressures by distinct abiotic and biotic environmental conditions above- and belowground and to different physicochemical properties such as volatility that are important in the air *versus* the rhizosphere [20]. When distinguishing between chemotypes, the respective chemical family as well as the organ should always be clearly specified [20], since chemotypes can vary depending on the organ, as evidenced in the present data for *C. hirsuta*.

For some accessions, quite contrasting patterns in richness and/or Shannon chemodiversity were found between leaves and roots. For example, plants from the one North American accession (US1) had the lowest Shannon diversity in leaf glucosinolates but among the highest in the roots, while plants of the European Dutch accession (NL1) showed an opposite pattern. Likewise, Shannon diversity differed between organs among accessions in different directions. Differences in leaf *versus* root richness and Shannon diversity of specialized metabolites among plants of different chemotypes have been described for a few plant species until now [76, 77] and may be a more widespread phenomenon, which is rarely investigated.

## Conclusion

Relative levels of four dominant glucosinolates in *Cardamine hirsuta* were found to be genetically controlled, using a comparison of plants grown at uniform conditions and comparing siblings. Observed relative levels suggested pent-4-enylglucosinolate (Peen) relative to but-3-enylglucosinolate (Buen) and phenethylglucosinolate (PE) relative to benzylglucosinolate (BZ) to be controlled by independent genetic polymorphic genes. Phenotypes varied markedly even among offspring from plants collected very close to each other, with local variation among plants collected a few km from each other apparently as large as variation among plants representing vast distances across Europe. Comparison with literature suggest relative levels of Peen relative to the sum of Buen and Peen to be due to a polymorphism in MAM genes, while the polymorphism in PE versus BZ seems to lack a previously described counterpart. Differences were also found in richness and Shannon diversity among the accessions, suggesting selection for high variation in glucosinolate profiles to be beneficial. Contrasting patterns between leaves and roots emphasize organ-specific biosynthesis and indicate that contrasting selection pressures must be at play above-and belowground. The study highlights that intraspecific chemodiversity needs to be investigated at various scales. The specific genetic underpinnings and ecological functions of this glucosinolate polymorphism in *C. hirsuta* have yet to be elucidated.

## List of abbreviations

HPLC-DAD: high performance liquid chromatograph coupled to diode-array detector
UHPLC-QToF-MS/MS: ultra-high performance liquid chromatograph coupled to quadrupole time of flight tandem mass spectrometer
GSL: glucosinolate
1mEt: 1-methylethylGSL
2mPr: 2-methylpropylGSL
1mPr: 1-methylpropylGSL
4mSOb: 4-(methylsulfinyl)butylGSL
5mSOp: 5-(methylsulfinyl)propylGSL
6mSOh: 6-(methylsulfinyl)hexylGSL
Buen: but-3-enylGSL
Peen: pent-4-enylGSL
BZ: benzylGSL
PE: phenethylGSL
4hBZ: 4-hydroxybenzylGSL
IM: indol-3-ylmethylGSL
1moIM: 1-methoxyindol-3-ylmethylGSL
4moIM: 4-methoxyindol-3-ylmethylGSL

## Declaration

### Ethics approval and consent to participate

Not applicable.

### Consent for publication

Not applicable.

### Availability of data and materials

The dataset of the glucosinolates for leaves and roots (absolute and relative composition) is provided in Supplement Table S1. The GenBank accession codes are provides in Supplement Table S2.

## Authors’ contributions

NA and CM conceived the ideas and proposed the conception of the experimental design. KD performed the glucosinolate analysis by HPLC-DAD, NA via UHPLC-QToF-MS/MS. MK performed barcoding. DZ performed the statistical analysis. NA and CM wrote the main sections of the manuscript and all authors reviewed the manuscript and gave final approval for publication.

## Supplementary Information

The online version contains supplementary material.

**Table S1**: Absolute and relative glucosinolate values of leaves and roots of offspring of 18 accessions of *Cardamine hirsuta*.

**Table S2**: GenBank accession codes of used *Cardamine hirsuta* material.

**Figure S1**: Distinction of 1-methylpropylglucosinolate (1mPr) and *n*-butylglucosinolate (Bu) in nine *Cardamine hirsuta* samples by UHPLC-QToF-MS of desulfated glucosinolates.

**Figure S2**: Examples of *m*/*z*, *t*_R_ and MS2 confirmation, in one selected sample each, of desulfated derivatives of all 14 identified glucosinolates are provided.

**Figure S3**: Levels of benzylglucosinolate (BZ) relative to the sum of BZ and phenethylglucosinolate (PE), and of pent-4-enylglucosinolate (Peen) relative to the sum of but-3-enylglucosinolate (Buen) and Peen when assuming the level of Peen to be LOD/2 = 0.05 µmol/g d.w. *versus* assuming zero values.

## Acknowledgements

We thank Miltos Tsiantis (MPI Plant Breeding, Cologn, Germany) and Martin Lysák (Masaryk University, Brno, Czech Republic) for providing us with seeds from several accessions. Sabrina Nufer is acknowledged for help in glucosinolate analysis.

