## Supplementary material for "Intraspecific glucosinolate chemodiversity in *Cardamine hirsuta*": TableS2_and_Supplement_Figures

**Table S1**: Absolute and relative glucosinolate values of leaves and roots of offspring of 18 accessions of *Cardamine hirsuta*. This data are found in a separate Excel Table.

**Supplement Table S2.** Herbarium codes for dried plant material (HEID, herbarium of Heidelberg University; Germany; for accession origin see Table 1 in main manuscript) and GenBank accession codes (QB).

**trnL intron:**

| | |
|---|---|
| **HEID505622** | **QB285281** |
| **HEID505623** | **QB285282** |
| **HEID505624** | **QB285283** |
| **HEID505625** | **QB285284** |
| **HEID505626** | **QB285285** |
| **HEID505627** | **QB285286** |
| **HEID505628** | **QB285287** |
| **HEID505629** | **QB285288** |
| **HEID505630** | **QB285289** |
| **HEID505631** | **QB285290** |
| **HEID505632** | **QB285291** |
| **HEID505633** | **QB285292** |
| **HEID505634** | **QB285293** |
| **HEID505635** | **QB285294** |
| **HEID505636** | **QB285295** |
| **HEID505637** | **QB285296** |
| **HEID505638** | **QB285297** |
| **HEID505639** | **QB285298** |
| **HEID505640** | **QB285299** |
| **HEID8013042** | **QB285300** |
| **HEID8013043** | **QB285301** |
| **HEID8013044** | **QB285302** |
| **HEID8013045** | **QB285303** |
| **HEID8013046** | **QB285304** |
| **HEID8013047** | **QB285305** |
| **HEID8013048** | **QB285306** |
| **HEID8013049** | **QB285307** |
| **HEID8013050** | **QB285308** |
| **HEID8013051** | **QB285309** |
| **HEID8013052** | **QB285310** |
| **HEID8013053** | **QB285311** |
| **HEID8013054** | **QB285312** |
| **HEID8013055** | **QB285313** |
| **HEID8013056** | **QB285314** |
| **HEID8013057** | **QB285315** |

**HEID8013058 QB285316**
**HEID8013059 QB285317**

**trnLF intergenic spacer:**
**HEID506622 QB285324**
**HEID506623 QB285325**
**HEID506624 QB285326**
**HEID506625 QB285327**
**HEID506626 QB285328**
**HEID506627 QB285329**
**HEID506628 QB285330**
**HEID506629 QB285331**
**HEID506630 QB285332**
**HEID506631 QB285333**
**HEID506632 QB285334**
**HEID506633 QB285335**
**HEID506634 QB285336**
**HEID506635 QB285337**
**HEID506636 QB285338**
**HEID506637 QB285339**
**HEID506638 QB285340**
**HEID506639 QB285341**
**HEID506640 QB285342**
**HEID8013042 QB285343**
**HEID8013043 QB285344**
**HEID8013044 QB285345**
**HEID8013045 QB285346**
**HEID8013046 QB285347**
**HEID8013047 QB285348**
**HEID8013048 QB285349**
**HEID8013049 QB285350**
**HEID8013050 QB285351**
**HEID8013051 QB285352**
**HEID8013052 QB285353**
**HEID8013053 QB285354**
**HEID8013054 QB285355**
**HEID8013055 QB285356**
**HEID8013056 QB285357**
**HEID8013057 QB285358**
**HEID8013058 QB285359**
**HEID8013059 QB285360**

**ITS:**

**HEID505622 QB269109**
**HEID505623 QB269110**
**HEID505624 QB269111**
**HEID505625 QB269112**
**HEID505626 QB269113**
**HEID505627 QB269114**

HEID505628 QB269115
HEID505629 QB269116
HEID505630 QB269117
HEID505631 QB269118
HEID505632 QB269119
HEID505633 QB269120
HEID505634 QB269121
HEID505635 QB269122
HEID505636 QB269123
HEID505637 QB269124
HEID505638 QB269125
HEID505639 QB269126
HEID505640 QB269127
HEID8013042 QB269142
HEID8013043 QB269143
HEID8013044 QB269144
HEID8013045 QB269145
HEID8013046 QB269146
HEID8013047 QB269147
HEID8013048 QB269148
HEID8013049 QB269149
HEID8013050 QB269150
HEID8013051 QB269151
HEID8013052 QB269152
HEID8013053 QB269153
HEID8013054 QB269154
HEID8013055 QB269155
HEID8013056 QB269156
HEID8013057 QB269157
HEID8013058 QB269158
HEID8013059 QB269159

**Figure S1A+B**. Distinction of 1-methylpropylglucosinolate (1mPr) and *n*-butylglucosinolate (Bu) in nine *Cardamine hirsuta* samples by UHPLC-QToF-MS of desulfated glucosinolates at a column temperature of 60 degrees {Leijonhufvud, under review #13191}. As controls, authentic references of Bu, 1mPr and 2-methylpropylglucosinolate (2mPr) were analyzed before samples and Bu and 1mPr after. A sample of desulfated glucosinolates from *Arabidopsis thaliana* Hi-0 seeds with established content of Bu and 2mPr {Leijonhufvud, under review #13191} was also included. 1mPr, not Bu, was also confirmed by MS2 (Fig. S2B).

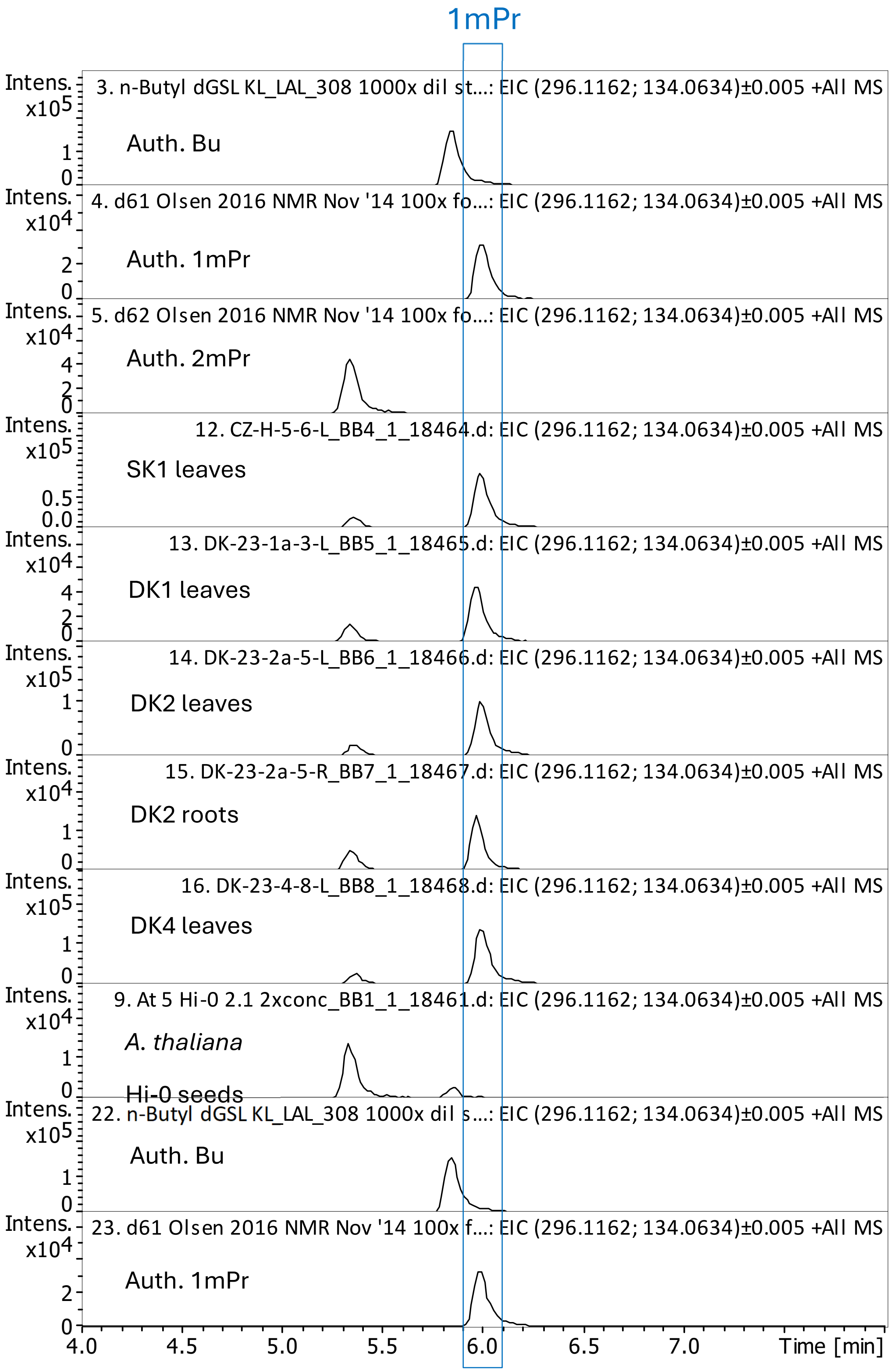

**Figure S1B**. Distinction of 1-methylpropylglucosinolate (1mPr) and *n*-butylglucosinolate (Bu) in four additional *Cardamine hirsuta* samples by UHPLC-QToF-MS of desulfated glucosinolates at a column temperature of 60 degrees {Leijonhufvud, under review #13191}. The same controls as in Fig. S1A were included.

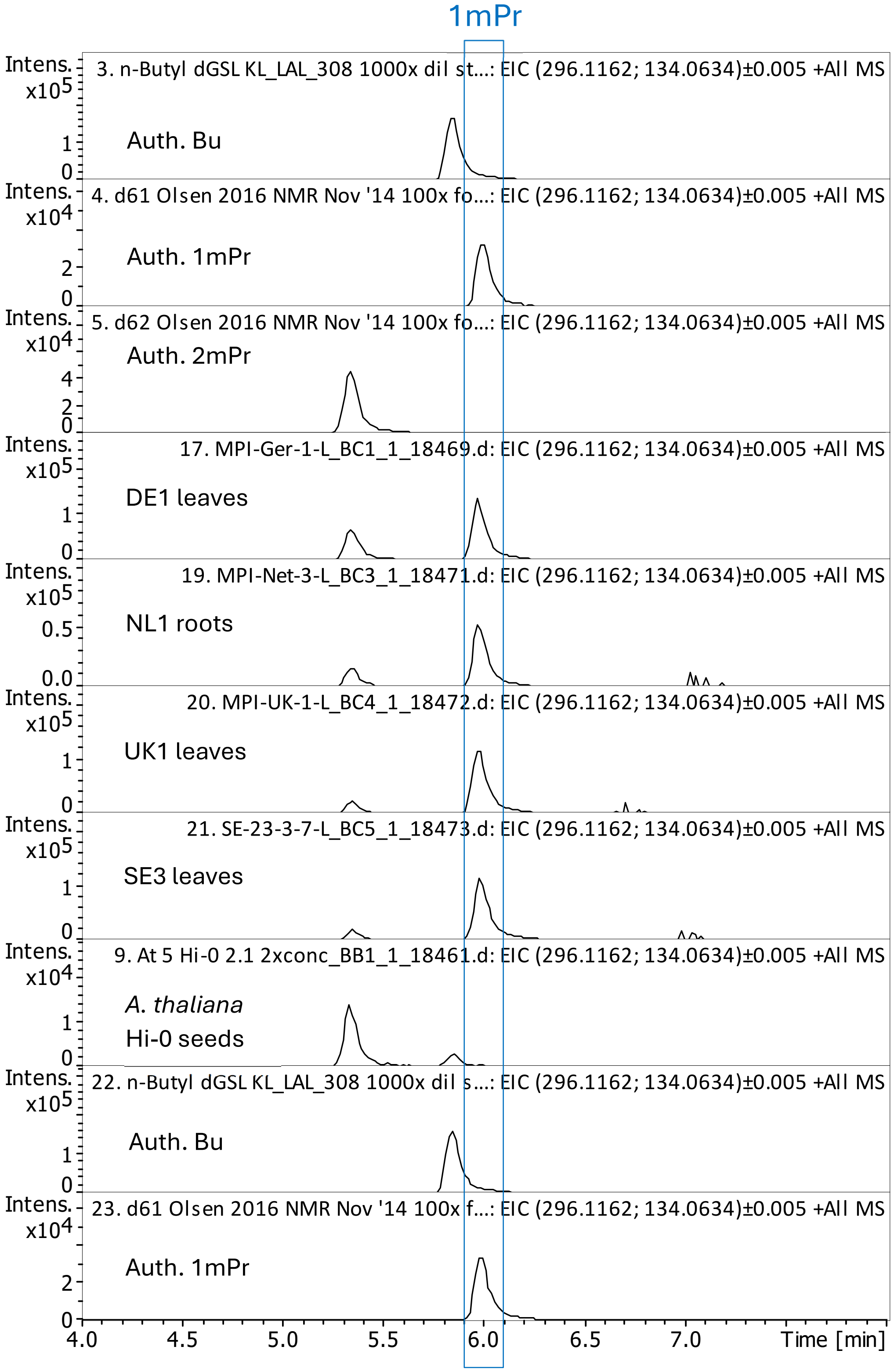

**Figure S2 A.** Examples, one plant sample each, of confirmation of correct *m*/*z*, $t_R$ and MS2 of desulfoglucosinolates (GSL) derivatives of the 14 GSL quantified by HPLC-DAD in the analysed samples. Peaks were first inspected in accession DE1 leaves. If MS2 not obtained in this sample, other accessions (as indicated in each case) were inspected with specific sample inspected guided by peak intensity.

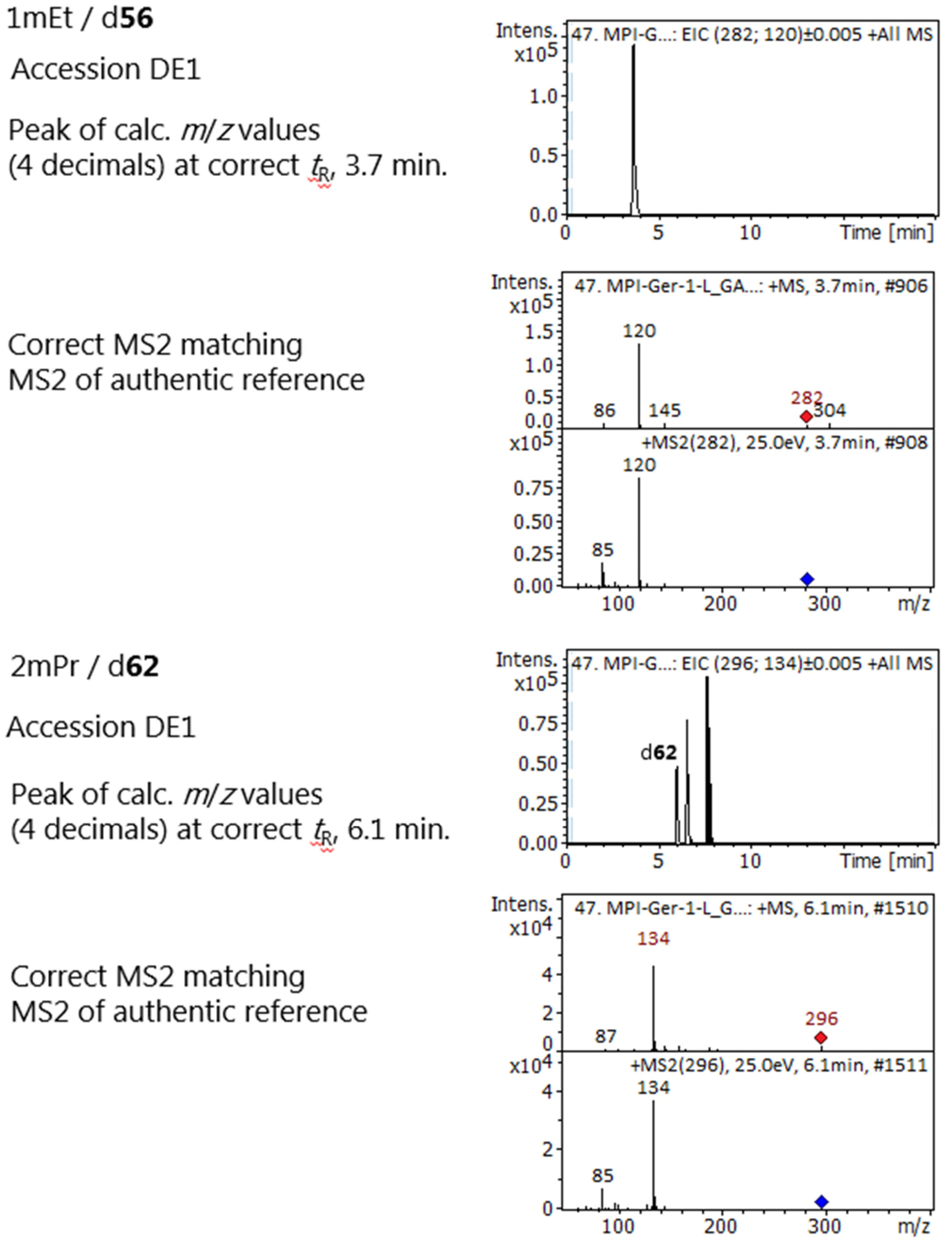

**Figure S2** (continued) **B.**

1mPr / d**61**

Accession DE1

Peak of calc. *m/z* values
(4 decimals) at correct $t_R$, 6.6 min.

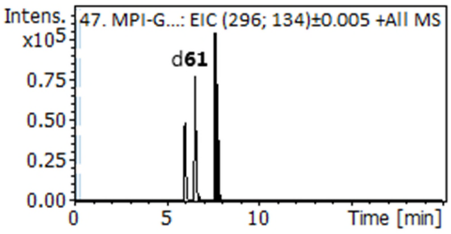


Correct MS2 matching
MS2 of authentic reference

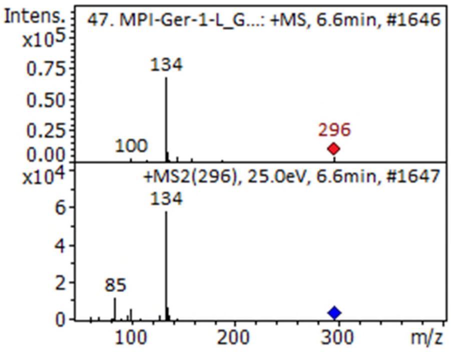


Coeluting Bu / d**[13]** has the same $t_R$ and MS2 of $[M+H]^+$,
so for distinction measured MS2 of $[M\text{-ahGlc}+H]^+$ fragment at *m/z* 134.
Dominance of *m/z* 83.07 rather than m/z 82.06 signifies that the peak
is mainly or entirely due to 1mPr / d**61**, not Bu / d**[13]** (Agerbirk et al., 2026). T
his conclusion was further supported by chromatographic separation at another
column temperature (Fig. S1)

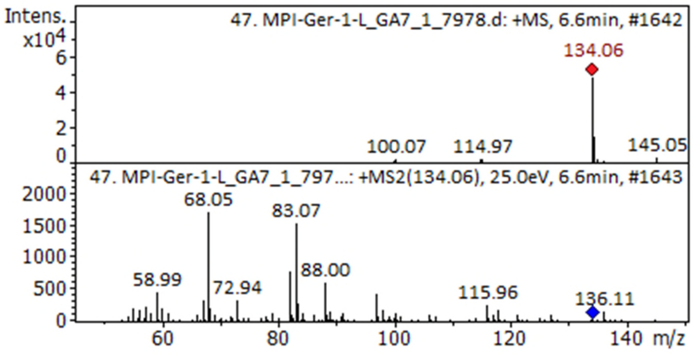

**Figure S2** (continued) **C.**

4mSOb / d**64**

Accession DE1

Peak of calc. *m/z* values
(4 decimals) at correct $t_R$, 1.8 min.

Correct MS2 matching
MS2 of authentic reference

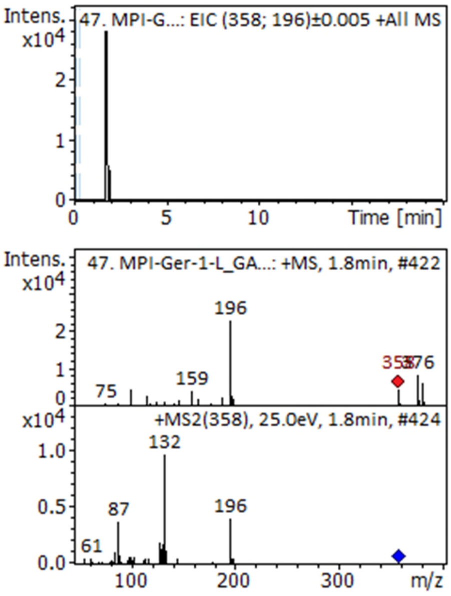


5mSOp / d**72**

Accession DE1

Peak of calc. *m/z* values
(4 decimals) at correct $t_R$, 3.6 min.
Compared to corresponding
peak from *A. thaliana* Col-0 seeds

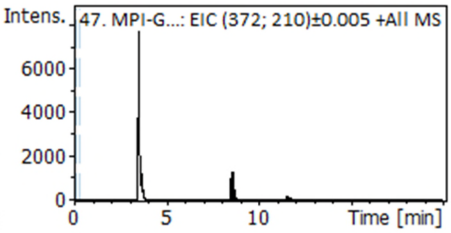


MS with $[M+H]^+$ but
MS2 not obtained
in this sample, see MS2 from
sample of SK1 next page

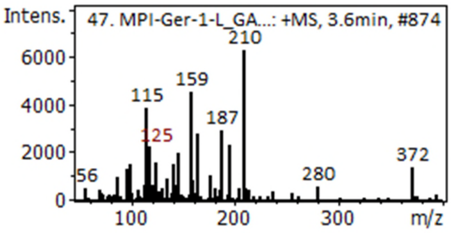

**Figure S2** (continued) **D.**

5mSOp / d**72**

Accession SK1

Peak of calc. *m/z* values (4 decimals) at correct $t_R$, 3.6 min. for the corresponding peak from *A. thaliana* Col-0 seeds

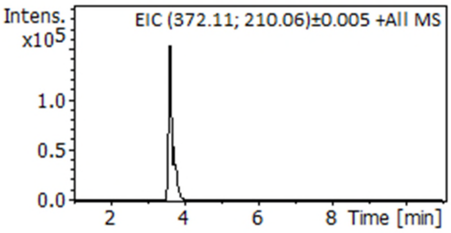


Correct MS2 compared to equivalent peak in *A. thaliana* Col-0 seed

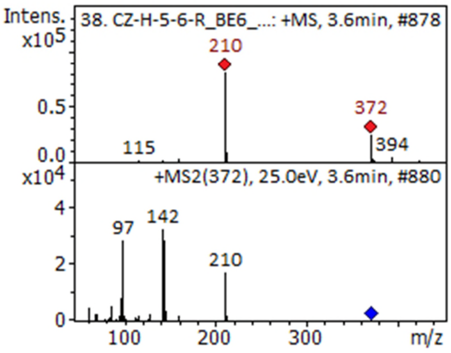


6mSOh / d**67**

Accession SK1

Peak of calc. *m/z* values (4 decimals) at correct $t_R$, 6.2 min. for the corresponding peak from other plant species such as *A. thaliana* Col-0 seeds and *Turritis glabra* leaves.

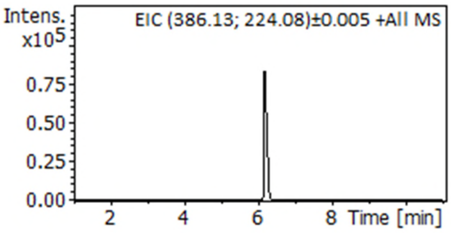


Correct MS2 compared to equivalent peak from *A. thaliana* Col-0 seed

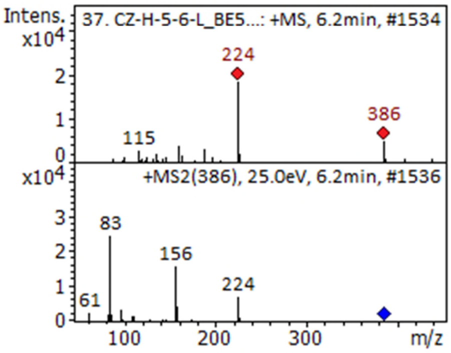

**Figure S2** (continued) **E.**

Buen / d**12**

Accession DE1

Peak of calc. *m/z* values
(4 decimals) at correct $t_R$, 4.9 min.

Correct MS2 matching
MS2 of authentic reference

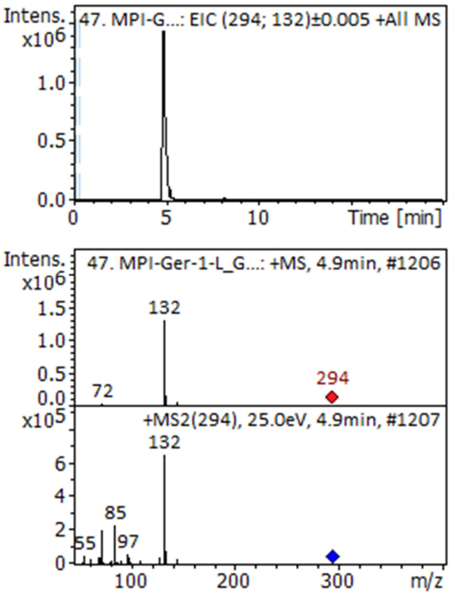


Peen / d**101**

Accession DE1

Peak of calc. *m/z* values
(4 decimals) at correct $t_R$, 7.5 min.

Correct MS2 matching
MS2 of authentic reference

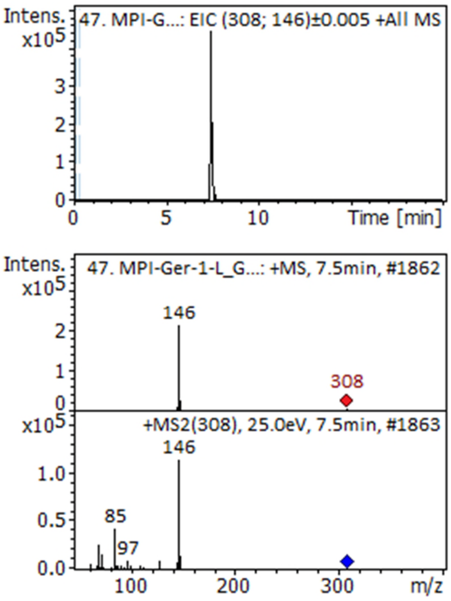

**Figure S2** (continued) **F.**

BZ / d**11**

Accession DE1

Peak of calc. *m/z* values at correct $t_R$, 7.7 min.

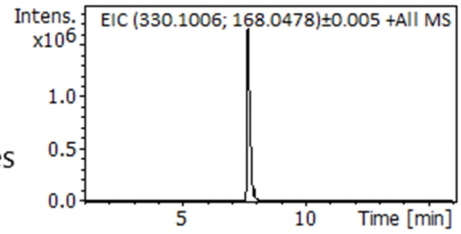


Correct MS2 matching authentic reference

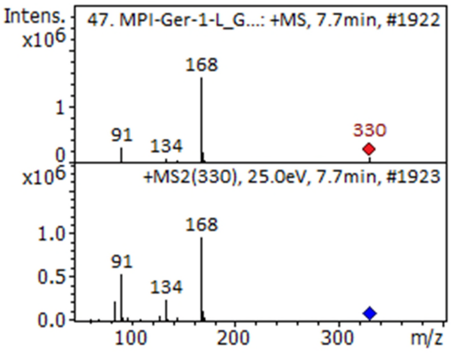


4hBZ / d**23**

Accession DK1 (DK-23-1a-3-L)

Peak of calc. *m/z* values (4 decimals) at correct $t_R$, 3.7 min.

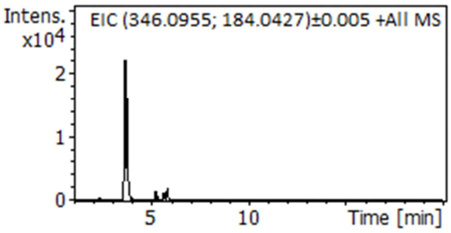


MS with $[M+H]^+$ but not MS2 not obtained in this sample. Confirmation of MS2 of *m/z* 184: next page

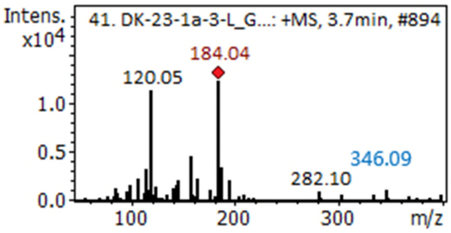

**Figure S2** (continued) **G.**

4hBZ / d**23** Accession DK1 (DK-23-1a-3-L)

Since MS2 of molecular proton adduct not obtained, *m/z* 184 (loss of anhydroGlc) fragment studied in MS2:

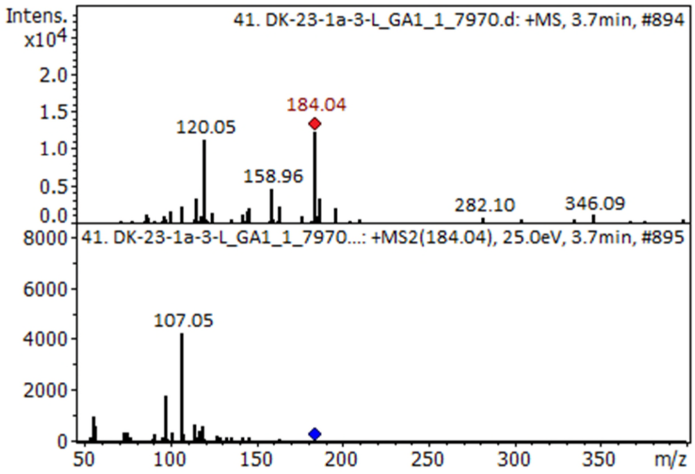


For comparison the corresponding MS2 of authentic d**23**:

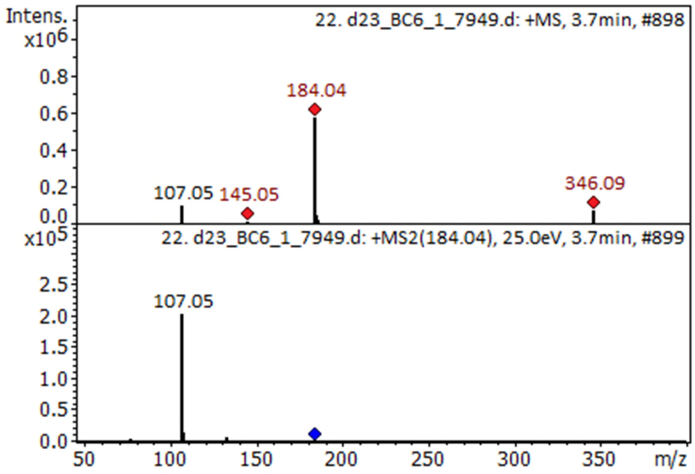


Conclusion: MS2 match sufficiently well for acceptance, since very minor MS2 spectra are known to sometimes show minor impurity ions

**Figure S2** (continued) **H.**

PE / d**105**

Accession DE1

Peak of calc. *m/z* values (4 decimals) at correct $t_R$, 9.9 min.

Correct MS2 matching MS2 of authentic reference

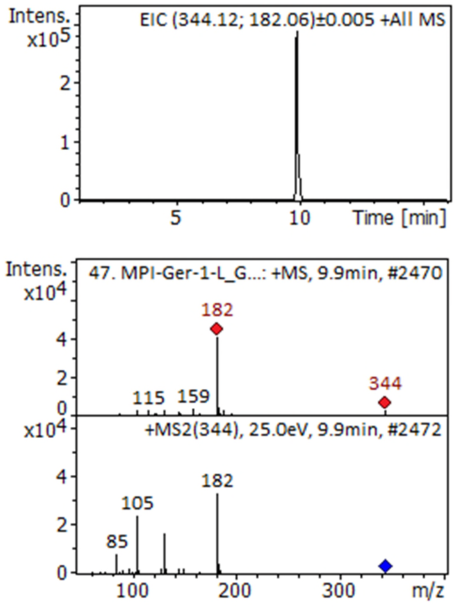


IM / d**43**

Accession DE1

Peak of calc. *m/z* values (4 decimals) at correct $t_R$, 8.6 min.

Correct MS2 matching MS2 of authentic reference

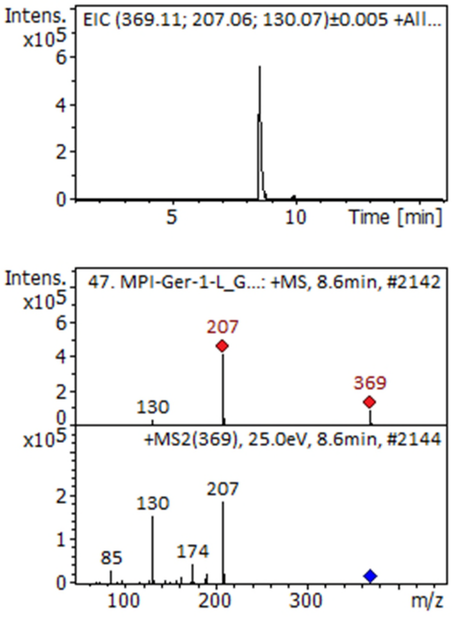

**Figure S2** (continued) **I.**

4moIM / d**48**

Accession DE1

Peak of calc. *m/z* values (4 decimals) at correct $t_R$, 9.9 min.

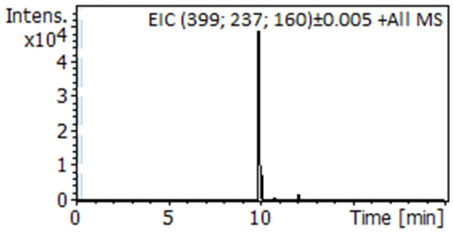


Correct MS2 matching MS2 of authentic reference

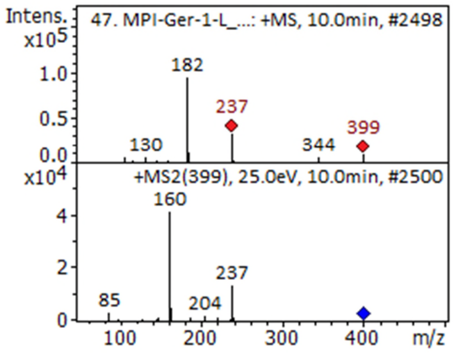


1moIM / d**47**

Accession DE1

Minor peak of calc. *m/z* values (4 decimals) at correct $t_R$, 12.1 min.

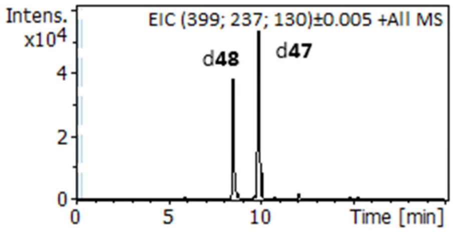


MS with $[M+H]^+$ but not MS2 not obtained in this sample (but see below for another sample)

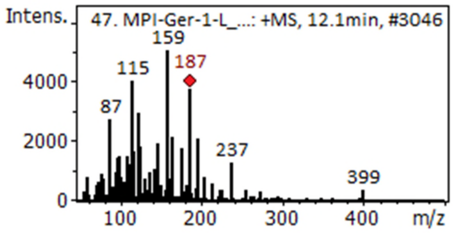

**Figure S2** (continued) **J.**

1moIM / d**47**

Accession GR1 (MPI-Gre-1-R): 1moIM / d**47**

Peak of calc. *m/z* values (4 decimals) at correct $t_R$, 12.1 min.

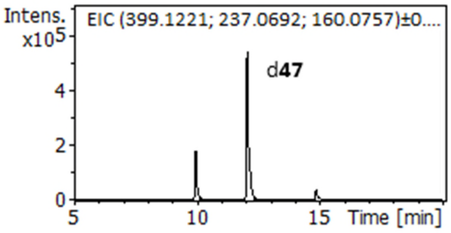


Correct MS2 matching MS2 of authentic reference

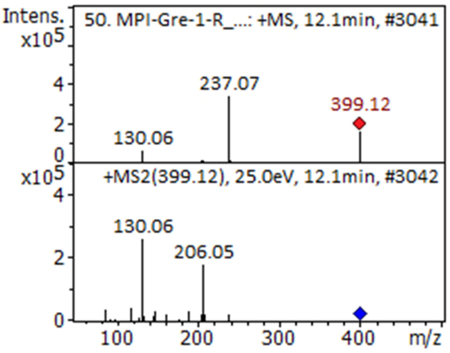

**Figure S3.** Levels of benzylglucosinolate (BZ) relative to the sum of BZ and phenethylglucosinolate (PE), and of pent-4-enylglucosinolate (Peen) relative to the sum of but-3-enylglucosinolate (Buen) and Peen.Consequence of assuming the level of Peen to be LOD/2 = 0.05 µmol/g rather than zero in 14 root samples for which this glucosinolate was not quantified by HPLC-DAD. Only one sample (red circle) swapped groups (from "low Peen" to "intermediate Peen" by being changed from plain zero Peen because of no quantification to LOD/2. The change brought this observation from being an outlier to being in agreement with several other samples from the same area of Sweden.

A: Plot using registered zero Peen levels directly for calculation of index

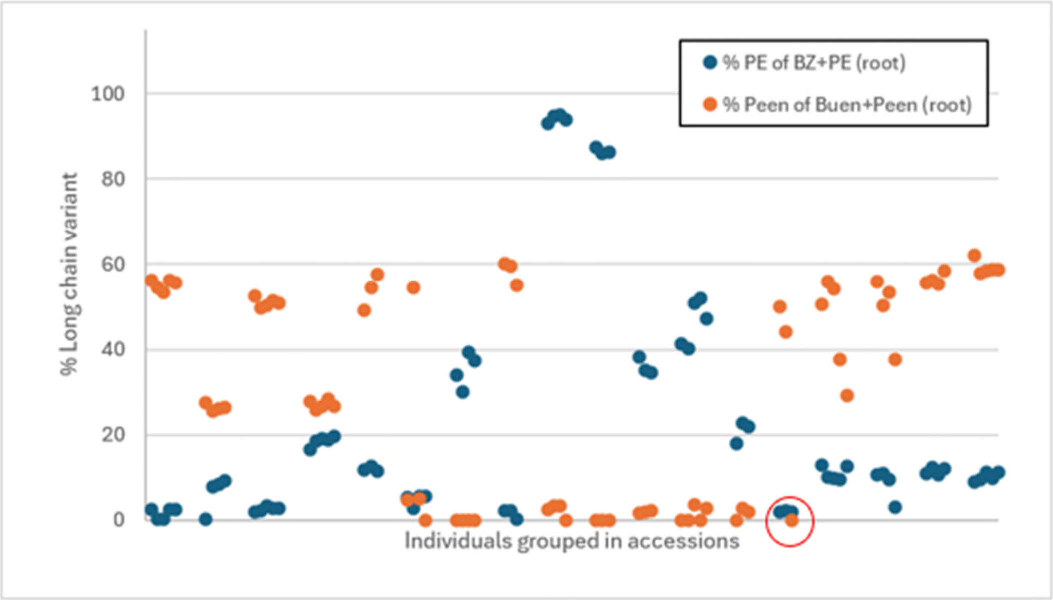


B: Plot after changing registered zero Peen levels to half estimated limit of detection

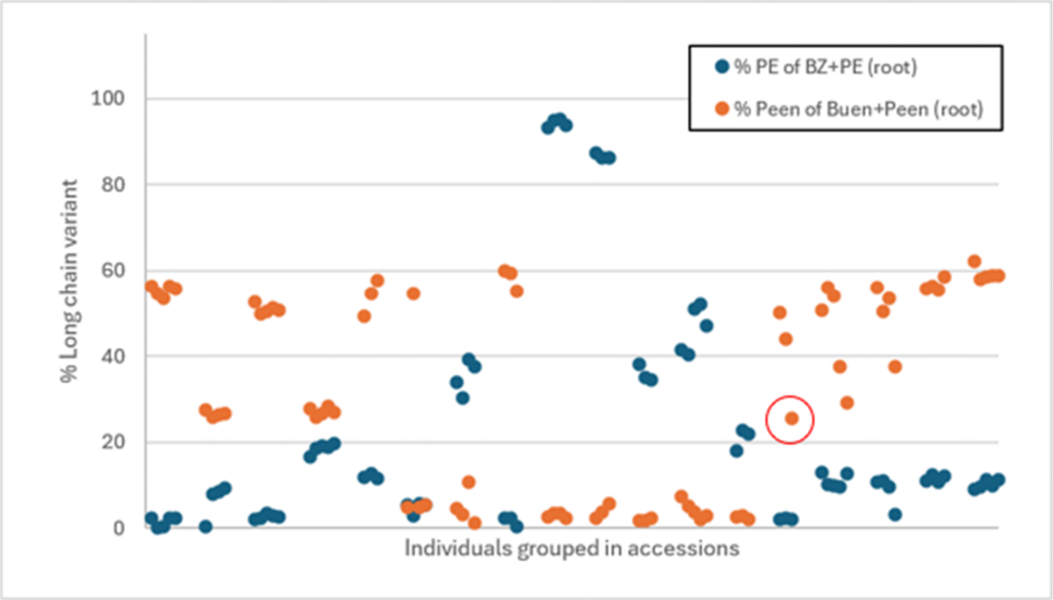
